# VP-CLEM-Kit: An accessible pipeline for visual proteomics using super resolution volume correlative light and electron microscopy (SR-vCLEM)

**DOI:** 10.1101/2025.04.08.647770

**Authors:** Dumisile Lumkwana, Jonathan Lightley, Arturo G. Vesga, Joost de Folter, Marie-Charlotte Domart, Catherine Maclachlan, Sunil Kumar, James Evans, Christopher Peddie, Alana Burrell, Azumi Yoshimura, Asandile Mangali, Nicola Vahrjmeijer, Meenal Bhaga, Catherine Smit, Benjamin Titze, Mathew Horrocks, Lorian Cobra Straker, Edwin Garcia, Martin Jones, Adam Mclean, Candice Roufosse, Ben Loos, Sonia Gandhi, Amy Strange, Ricardo Henriques, Paul French, Lucy Collinson

**Author notes:** These authors contributed equally to the work (Dumisile Lumkwana, Jonathan Lightley, Arturo G. Vesga, Joost de Folter). Corresponding authors. Corresponding author email addresses.

## Abstract

Visual proteomics (VP) aims to allow researchers to visualise, measure and analyse proteins in the context of cell and tissue structure in health and disease. VP is becoming a reality through technological advances across several domains, including *in situ* structural biology and correlative light and electron microscopy (CLEM). However, widespread adoption remains limited due to the complexity and cost of the various VP approaches reported to date. Here we present the VP-CLEM-Kit, a disruptive cost-effective pipeline for super resolution volume CLEM (SR-vCLEM) that can be implemented with minimal advanced electron microscopy expertise and equipment, making it accessible to light microscopy facilities and research labs. SR-vCLEM is based on in-resin fluorescence (IRF), where fluorophores are preserved through processing into resin. The easyIRF protocol reported here reduces the requirement for complex costly sample preparation equipment and toxic chemicals compared to standard IRF protocols. easyIRF blocks are cut into ultrathin sections that are imaged using ‘tomoSTORM’, a new modular and cost-effective *openFrame*-based light microscope controlled by the open-source software package Micro-Manager that provides serial single molecule localisation microscopy in array tomography format. Sections are then post-stained and imaged using a tabletop scanning electron microscope controlled by open-source SBEMimage software to run in array tomography format. We demonstrate the potential of the VP-CLEM-Kit by imaging organelle reporters in human cell lines and stem cell derived neurons, fluorescently labelled protein in neurons, and immunolabelled cells in human kidney biopsy tissue from transplant patients. The VP-CLEM-Kit delivers a ∼5-fold improvement in resolution at a ∼7-fold lower cost, and thus provides new technical capability as well as a blueprint for more equitable access to advanced imaging workflows.

## Introduction

Huge progress has been made in mapping the three-dimensional structure of macromolecules using cryo-electron microscopy (cryoEM) (Ignatiou et al., 2024), and of cells and tissues using volume electron microscopy (volume EM; vEM) (Peddie et al., 2022). There is now a concerted effort by different communities to map the spatial distribution of macromolecules in the context of cell and tissue structure (Hutchings and Villa, 2024; McCafferty et al., 2024). These maps will underpin a new understanding of the function of molecules in biological context in health and disease.

There are several different approaches for spatial mapping of macromolecules in the context of cell and tissue structure, each operating at different scales and resolutions, and collectively referred to as visual proteomics (VP) (Nickell et al., 2006). In situ structural biology (ISSB) images macromolecules at near-atomic resolution in the context of near-native-state cell membranes using cryo-electron tomography and subtomogram averaging (Briggs, 2013). Volume correlative light and electron microscopy (vCLEM) uses fluorescent tags to determine the spatial localisation of macromolecules through larger cell and tissue volumes, followed by preparation of the same sample for electron imaging and merging of the two image datasets (Guerin and Lippens, 2021). Each technique has limitations. ISSB is limited by the penetration of electrons through organic material to a continuous volume of ∼300 nm, and vCLEM uses fluorophores as a proxy for macromolecules as it lacks the sample preparation and resolution to image the macromolecular structures themselves. A further limitation of both techniques, in terms of application to real-world research, is the cost and complexity of the workflows and equipment required, resulting in VP being available to only a handful of well-funded specialist research groups and core facilities. The VP-CLEM-Kit pipeline aims to break through this access barrier, whilst also delivering improvements in the resolution and precision of vCLEM.

To date, vCLEM has generally been performed using a pre-embedding CLEM protocol (Polishchuk et al., 2000). Cells or tissues expressing fluorescently-tagged markers are first fixed using 4% (para)formaldehyde and then imaged using fluorescence microscopy. The samples are then prepared for EM with: a secondary fixation using formaldehyde and glutaraldehyde to crosslink proteins; heavy metal staining to crosslink lipids and add contrast to the membranes; dehydration to remove water; infiltration with a liquid epoxy resin; and polymerisation of the resin using heat to form a hard block. The block can then be sectioned and imaged using a range of volume EM technologies depending on the field of view (FOV) and resolution required. Volume EM options (Peddie et al., 2022) include serial section transmission EM (ssTEM) (Harris et al., 1992; White et al., 1994; White et al., 1986), serial section electron tomography (ssET) (Soto et al., 1994), GridTape® TEM (Graham et al., 2019; Yin et al., 2020), array tomography (AT) in a scanning EM (SEM) (Micheva and Smith, 2007), serial block face SEM (SBF-SEM) (Denk and Horstmann, 2004; Genoud et al., 2004), focused ion beam SEM (FIB-SEM) (Knott et al., 2008), enhanced FIB-SEM (eFIB) (Xu et al., 2017) or plasma FIB-SEM (pFIB) (Sergey et al., 2018). The fluorescence images are then overlaid onto the electron images to determine the identity of cells in the EM images or to localise the fluorescently-tagged molecule to subcellular structures to reveal information about their function in health and disease.

The advantages of pre-embedding vCLEM include the ability to choose the optimal fluorescence and electron microscope modalities for the biological sample and the research question, as well as optimal sample preparations for both light and electron microscopy because the two modalities are used independently at different stages of the workflow. However, there is a significant compromise in the accuracy of the overlay of fluorescence and electron images due to shrinkage and warping artefacts during sample preparation, and the difference in resolution and contrast type in the images. This limits the correlation precision to approximately 1 µm.

The last decade has seen the introduction of in-resin fluorescence (IRF) techniques (Heiligenstein and Lucas, 2022) that aim to overcome the limited correlation precision of pre-embedding vCLEM techniques, particularly for VP experiments where protein localisation to structure is the aim. Samples are fixed using a high-pressure freezer, freeze substituted in a solvent with low percentages of water and uranyl acetate, and embedded in methacrylate resins that are polymerised using UV radiation. The result is preservation of fluorescent proteins and dyes in the resin block that, when cut into ultrathin sections, can be imaged first in a fluorescence microscope and then in an electron microscope. The overlay accuracy of the LM to EM images can thus be improved by an order of magnitude (from ∼1 micron to ∼100 nm) (Kukulski et al., 2011). This is because the resolution of the fluorescence images is closer to that of the EM images. The axial resolution is determined by the thickness of the section (usually well below the diffraction limit at 50 – 200 nm), noting that out of plane fluorescence is not present for thin sections. The lateral resolution can be further improved using single molecule localisation microscopy (SMLM), leveraging the observation that fluorescent proteins and dyes blink in-resin (Fu et al., 2020; Johnson and Kaufmann, 2017; Paez-Segala et al., 2015; Peddie et al., 2017).

Preservation of fluorescence in-resin comes at the expense of contrast in the electron images due to replacement of osmium tetroxide with uranyl acetate in the IRF sample preparation to preserve the fluorescence. However, even low percentages of uranyl acetate still appear to compromise the fluorescence signal to some extent. This can be observed in cell pellets embedded using the IRF technique that exhibit a gradient of electron contrast across the pellet (high contrast on the side exposed to the freeze substitution solution and low contrast on the side facing the metal planchette in which the pellet was frozen), associated with an opposing gradient in fluorescence preservation (Peddie et al., 2014). A number of new fluorescent probes have been developed that report some level of fluorescence preservation in the presence of osmium (Fu et al., 2020; Paez-Segala et al., 2015; Peng et al., 2022; Tanida et al., 2020a). However, preservation is generally reported to be ∼10-60% of the original fluorescence emission in the sample, so their use in IRF experiments generally requires biological systems in which the labelled protein has a high endogenous expression level or where overexpression does not alter the process being studied. A fluorescence preservation level of 10-60% also compromises the use of SMLM to increase lateral resolution, since the localisation precision depends on the number of photons detected per fluorophore, and fewer fluorophores would result in sparser labelling.

This work reports a new super resolution volume CLEM (SR-vCLEM) pipeline called the VP-CLEM-Kit, which was developed to improve the resolution and imaging volume of IRF, whilst reducing the expense and complexity of performing vCLEM. The VP-CLEM-Kit thus supports broader adoption of vCLEM in light microscopy facilities, research labs and lower resource settings. Given the challenges associated with equipment access, know-how and human capacity in such settings in general, and their limited exposure to volume EM and SMLM specifically, the VP-CLEM-Kit represents a new model to widen access to advanced imaging workflows.

## Results

### The VP-CLEM-Kit concept

This first version of the VP-CLEM-Kit provides a cost-effective, user-friendly pipeline for SR-vCLEM, combining a new sample preparation protocol with open-source and commercial imaging hardware, and open-source software solutions for hardware control and image analysis (Fig.1). The easyIRF protocol preserves fluorescent markers in cells and tissues embedded in resin, without the need for a high-pressure freezer and with minimal toxic chemicals. The sample is then cut into ultrathin sections, and the sections are picked up onto a conductive glass substrate. The tomoSTORM light microscope is an implementation of the cost-effective modular o*penFrame*-based microscopy system customised for serial SMLM of easyIRF sections in array tomography format. Existing open-source software (Picasso) (Schnitzbauer et al., 2017) has been leveraged for reconstruction of the SMLM images. A new open-source napari (Chiu et al., 2022) software plug-in named Microscopy Array Section Setup (MASS) has been designed to support the user to track regions of interest (ROIs) from the fluorescence microscope to the electron microscope. The commercial Phenom Pharos tabletop SEM (Thermo Fisher Scientific) is controlled with a customised version of the open-source software SBEMimage (Titze et al., 2018) to run in array tomography format. Serial LM and EM images are aligned using either the open-source Fiji (Schindelin et al., 2015) plug-in TrakEM2 (Cardona et al., 2012) or a new automated registration algorithm. Finally, LM and EM images are overlaid using the open-source Fiji plug-in BigWarp (Bogovic et al., 2016). The resulting overlays can then be used to assign molecular distributions to subcellular structures. Each component of the VP-CLEM-Kit workflow can be used independently, e.g. to generate volume SMLM data or volume EM data, or combined into the full SR-vCLEM pipeline.

**Figure 1.**
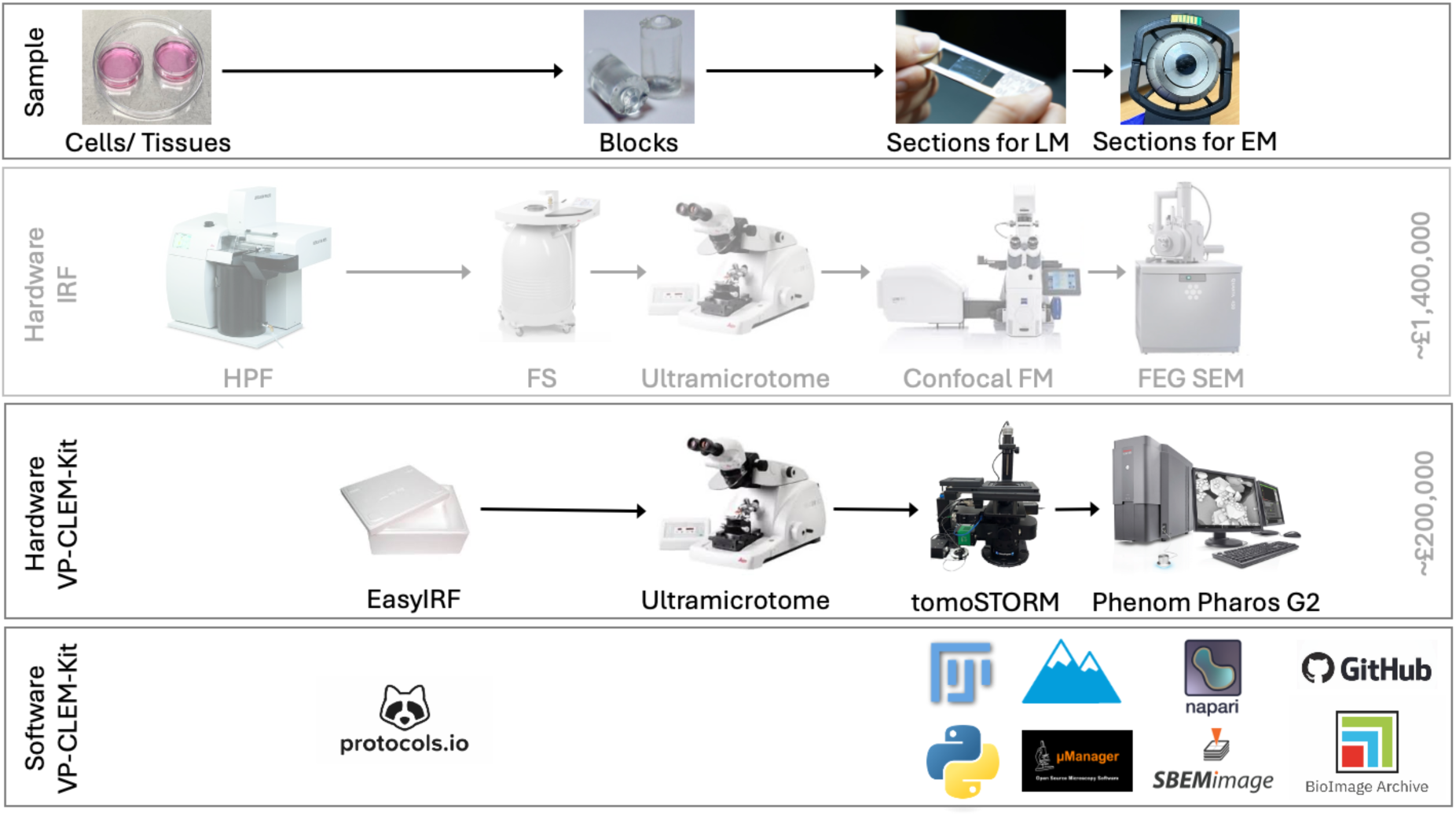
The VP-CLEM-Kit pipeline. The biological sample (illustrated by cells growing in culture) was fixed and embedded in resin using the easyIRF protocol. Fluorescent proteins and dyes were preserved. The sample was then cut into a series of ultrathin sections and the sections retrieved onto a conductive glass coverslip. The sections were imaged using a fluorescence microscope and then post-stained and imaged using an electron microscope. The VP-CLEM-Kit pipeline removes expensive complex sample preparation equipment (HPF and AFS), adds new functionality with the low-cost tomoSTORM light microscope, replaces the large floor-standing SEM with a tabletop version modified to run in array tomography mode, and leverages a suite of open-source software with new plug-ins and modes for hardware control and image analysis. The cost of the VP-CLEM-Kit pipeline (including a commercial ultramicrotome, which is currently required for cutting sections) is 7-fold less than the previous IRF pipeline. The kit can be installed in a light microscopy facility or research laboratory without the need for specialist environmental protection to mitigate vibration and electromagnetic fields.

### The easyIRF protocol

The easyIRF protocol (dx.doi.org/10.17504/protocols.io.3byl498x8go5/v1) was developed to reduce the cost and complexity of preparing IRF samples whilst maximising fluorescence preservation and fluorophore blinking, and introducing sufficient electron contrast to add ultrastructural information for CLEM.

To achieve this, the high-pressure freezer was removed from the workflow, and instead the samples were chemically fixed at 4℃ using 4% EM-grade (para)formaldehyde. After fixation, the samples underwent dehydration by progressive lowering of temperature (PLT). PLT helps to stabilise the sample by reducing the temperature of the sample (from 4 ℃ to -20 ℃) as the solvent concentration increases. Unlike most freeze substitution protocols, the easyIRF PLT medium does not contain any heavy metal stain, which maximises fluorescence preservation for downstream SMLM. It should be noted that the lack of osmium reduces crosslinking of lipids in the membranes, leading to a compromise in the downstream ultrastructural preservation. Once fully dehydrated, the sample was infiltrated at -20℃ with the methacrylate resin LR White and then polymerised at -20℃ using UV light. The entire protocol took 5 days from fixation to polymerised resin block, with around 6 hours of hands-on wet lab work required during that period.

Several studies report on the stability of fluorescence in IRF samples (Baatsen et al., 2021; Campbell et al., 2022; Heiligenstein and Lucas, 2022), with most studies showing that fluorescence reduces drastically over time. In easyIRF blocks, similar to our HPF IRF protocol (Peddie et al., 2014), fluorescence is maintained for at least 2 years (the age of the oldest samples at the time of publication).

In terms of resources, the easyIRF protocol requires a fridge and a fume hood, as well as some standard lab consumables (Table 1). The PLT can be performed in an automated freeze substitution unit (AFS), which uses a liquid nitrogen dewar to accurately control the temperature of the samples during the dehydration, resin infiltration and resin polymerisation steps. In labs or core facilities where an AFS and liquid nitrogen are not available, the dehydration and resin infiltration process can be performed in a -20℃ freezer or a walk-in -20℃ room. The resin polymerisation can be performed in a -20℃ freezer or a polystyrene box in a walk-in -20℃ room equipped with a UV torch.

**Table 1.**
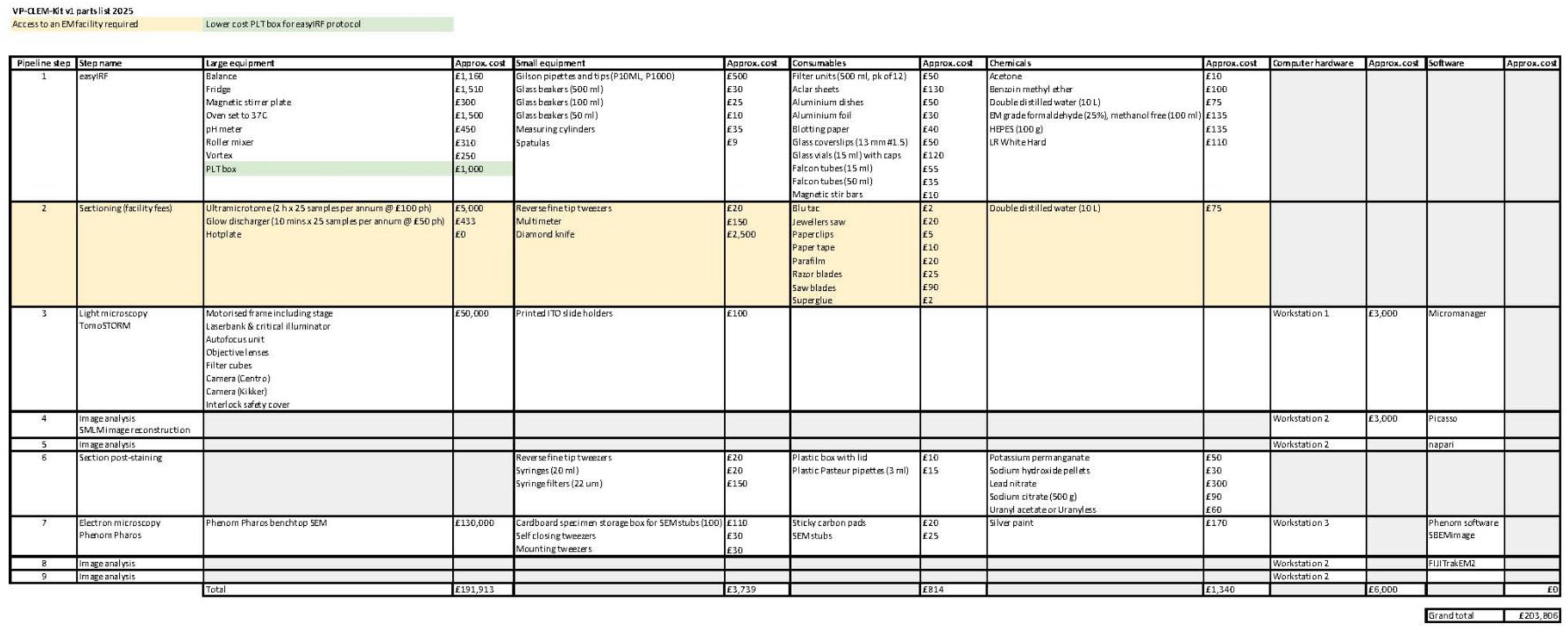

### Ultrathin sectioning of easyIRF blocks

The VP-CLEM-Kit currently requires access to an ultramicrotome with a diamond knife to cut ultrathin sections from easyIRF blocks for imaging in array tomography format. Consequently, the pipeline does depend on access to ultramicrotomy capability. For deployment of the VP-CLEM-Kit in a light microscopy facility or research lab, costs should be included for access to ultramicrotomy in a local or remote core facility or research lab (Table 1) or for the purchase of an ultramicrotome and training of a local specialist.

easyIRF sections were cut and retrieved onto conductive indium tin oxide (ITO)-coated glass coverslips for imaging. Sections were generally imaged for SMLM within 48 hours of sectioning to minimise the risk of a reduction in fluorescence over time, though fluorescence was visible more than a week after sectioning on coverslips that were stored at 4℃ in a fridge in the dark.

### Biological systems tested with the easyIRF protocol

The easyIRF protocol has been demonstrated to work for two formats of biological sample commonly analysed in correlative workflows - monolayers of cells grown on glass coverslips (Fig.2 & Fig.3) and tissue sections (Fig.4). The protocol successfully preserved green fluorescent protein (GFP), Hoechst, an organelle tracker dye (lysotracker), and Alexa Fluor dyes (AF488 and AF568) in these samples.

**Figure 2.**
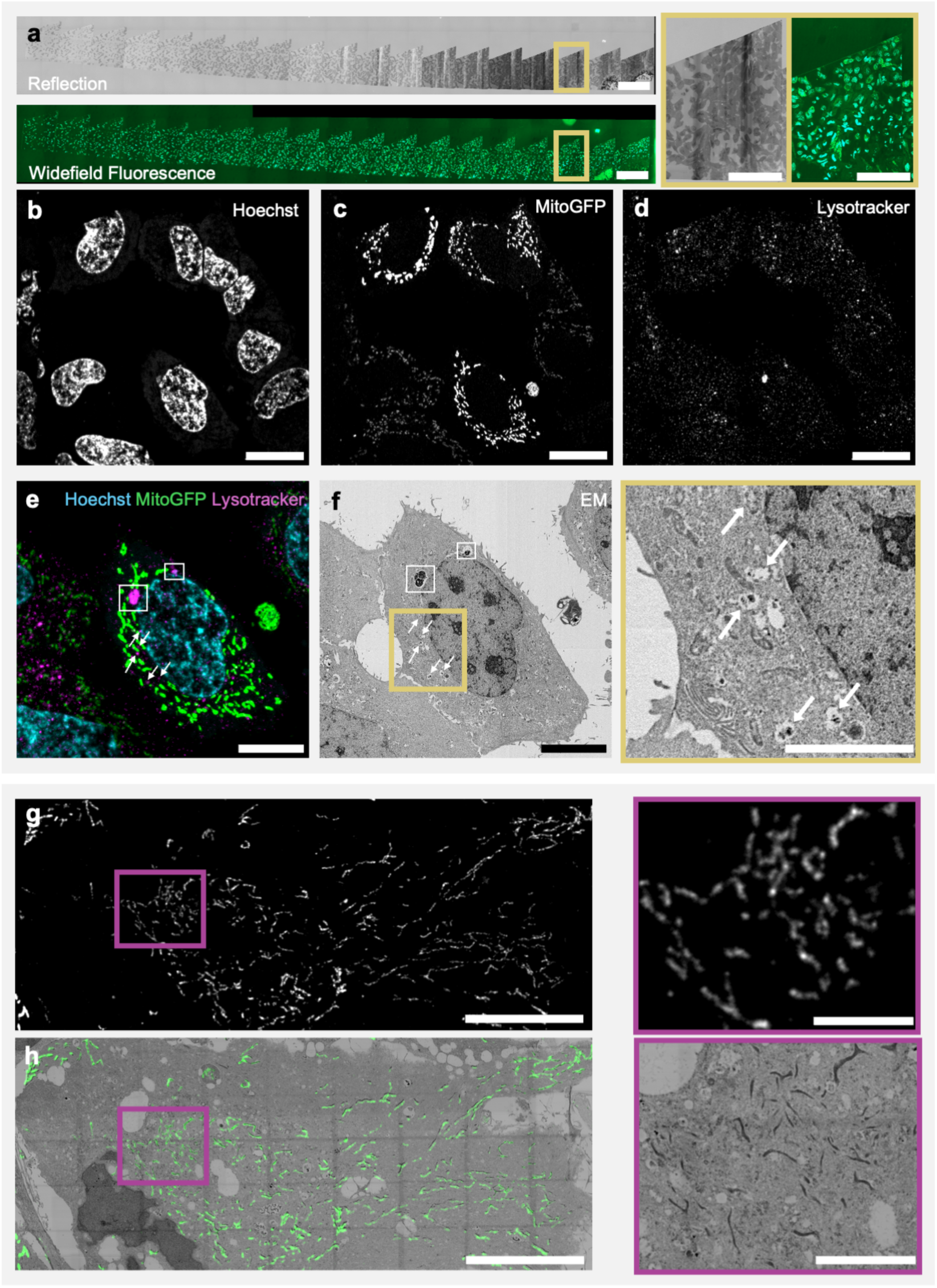
In-resin fluorescence and electron contrast in HeLa cells and hiPSC-derived cortical neurons expressing mitoGFP using the easyIRF protocol. (a) Overview RF and WF images showing serial sections of HeLa cells (scale bar: 400 µm) with insets showing zooms of one of the sections in each modality (scale bar: 200 µm). (b, c, d) High resolution LSM900 Airyscan images showing the preservation of Hoechst (nucleus), mitoGFP (mitochondria) and lysotracker (lysosomes) in a 200 nm thick easyIRF section (scale bar: 20 µm). (e) Cell of interest showing overlay of Hoechst, mitoGFP and lysotracker signals and (f) ultrastructure (scale bar: 10 µm), with inset showing preservation of mitochondria, nucleus, nuclear envelope, and large (white boxes) and small (arrows) lysosomes (scale bar: 2 µm). (g) hiPSC-derived cortical neurons showing preservation of mitoGFP in a 200 nm thick easyIRF section, and (h) the matching ultrastructure (scale bar: 20 µm). Insets show that the pattern of mitoGFP in the LM images matches the pattern of mitochondria in the EM image (scale bar: 5 µm).

**Figure 3.**
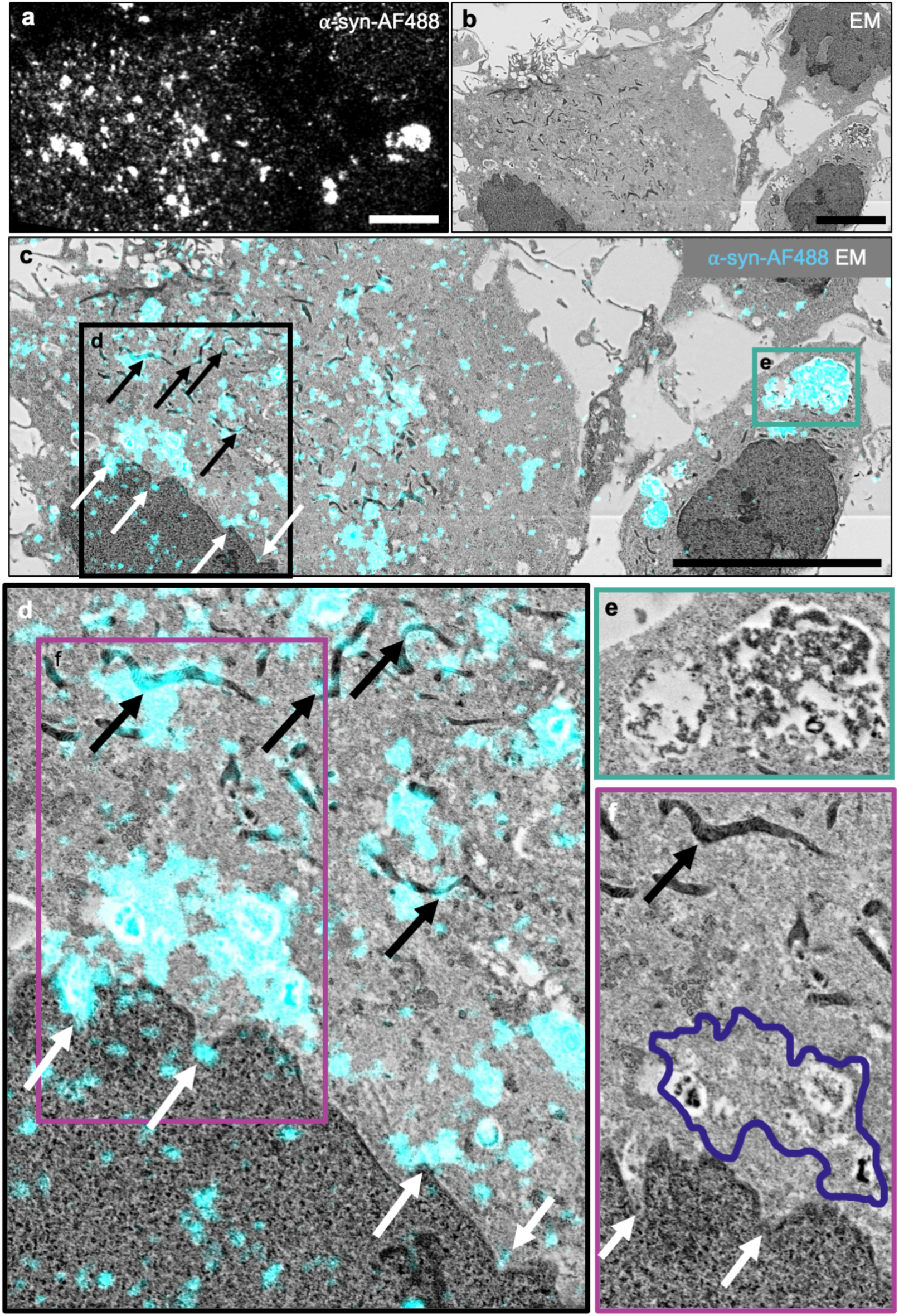
In-resin fluorescence and electron contrast in hiPSC-derived cortical neurons labelled with AF488-ɑ-Syn using the easyIRF protocol. (a) LSM900 Airyscan image showing preservation of AF488-ɑ-Syn in a 200 nm thick easyIRF section (scale bar: 20 µm). (b) EM image of the same region showing the ultrastructure of the cells. (c) A cell of interest showing the ɑ-Syn-AF488 fluorescence image overlaid onto the electron image from the same section (scale bar: 20 µm). (d) Higher magnification image of the black boxed region from panel c showing localisation of AF488-ɑ-Syn to mitochondria (black arrows), tortuous nuclear envelope foldings (white arrows) and lysosome-like structures. (e) Higher magnification image of the teal boxed region from panel c showing large lysosome-like structures filled with AF488-ɑ-Syn and electron-dense material. (f) Higher magnification image of the magenta boxed region from panel d showing the ultrastructure of the mitochondria, nuclear envelope and lysosome-like structures associated with AF488-ɑ-Syn labelling.

**Figure 4.**
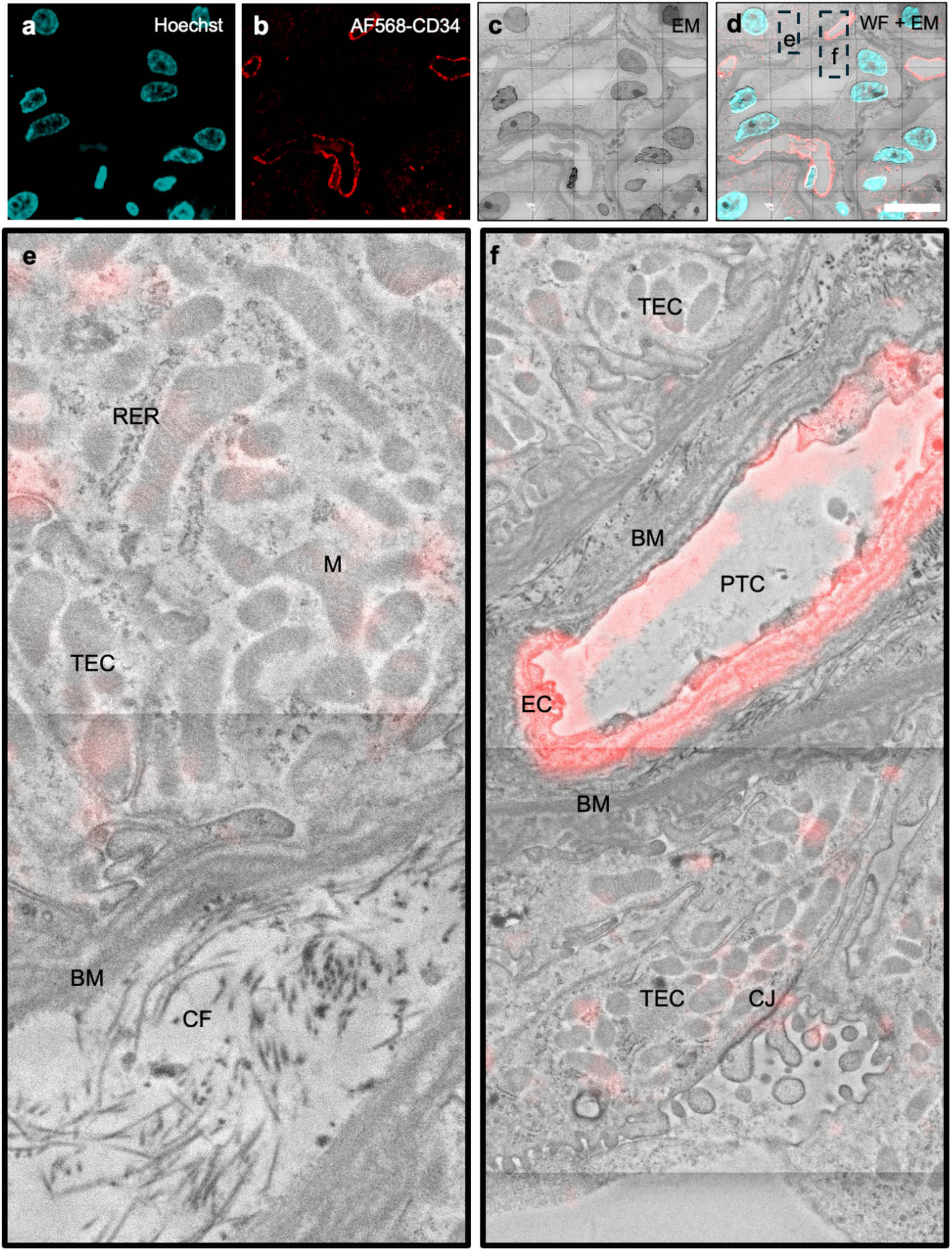
In-resin fluorescence and electron contrast in human kidney biopsy tissue immunolabelled with an anti-CD34 antibody and AF568 using the easyIRF protocol. (a) LSM900 Airyscan image showing preservation of Hoechst and (b) AF568-CD34 in a 200 nm thick easyIRF section (scale bar: 20 µm). (c) EM image of the same region showing the ultrastructure of the tissue. (d) Overlay of the LM and EM images. (e) Higher magnification image of a region from panel d showing ultrastructure of the kidney. (f) Higher magnification image of a region from panel d highlighting the endothelial cells of a peritubular capillary (PTC) identified by the CD34 label. RER - rough endoplasmic reticulum; M - mitochondria; CF - collagen fibrils; BM - basement membrane; CJ - cell junctions; TECs - tubular epithelial cells.

HeLa cells expressing mitoGFP for mitochondria and labelled with Hoechst for nuclei and lysotracker for lysosomes were used as a benchmark sample for protocol development (Fig.2). Human induced pluripotent stem cell (hiPSC)-derived cortical neurons expressing mitoGFP were used to test the protocol on a complex cell model that can recapitulate human central nervous system cell types *in vitro* (Fig.2). Successful application of the method to iPSC-derived cortical neurons opened the avenue to addressing a central nervous system (CNS) disease-related question, namely, whether it was possible to track the protein alpha-synuclein inside neurons. Alpha-synuclein (ɑ-Syn) is an intrinsically disordered protein in its monomeric state, but in Parkinson’s disease it can undergo self assembly and aggregate to form oligomeric and fibrillar species that form the pathological hallmark of the disease. The protocol was thus applied to hiPSC-derived cortical neurons exposed to 500nM of recombinant ɑ-Syn monomers conjugated to AF488 (ɑ-Syn-AF488) (Fig.3). This cell system also allowed comparison of correlation precision of the VP-CLEM-Kit with the same experiment performed with pre-embedding vCLEM (Choi et al., 2022), in which ɑ-Syn oligomers were localised to organelles with a precision of ∼1 micron. The easyIRF protocol was then tested on vibratome slices from a human kidney biopsy labelled with an anti-CD34 primary antibody and a secondary antibody conjugated to AF568, to locate endothelial cells in the glomeruli and peritubular capillaries in studies of antibody-mediated kidney transplant rejection (Fig.4).

### Imaging easyIRF sections in commercial light and electron microscopes

A commercial confocal light microscope (Zeiss LSM900 Airyscan) and a commercial floor-standing SEM (Thermo Fisher Scientific Quanta FEG SEM or Zeiss GeminiSEM) were used to image easyIRF sections during the development phase of the easyIRF protocol (Fig.2). The ITO-coated coverslips were first imaged in reflection (RF) mode to find the sections and then in widefield fluorescence (WF) mode to find fluorescent regions within each section (Fig.2a). Hoechst (Fig.2b), mitoGFP (Fig.2c) and lysotracker (Fig.2d) were successfully preserved and imaged in 200 nm thick sections of easyIRF-prepared HeLa cells using the Airyscan mode.

After fluorescence microscopy, sections were post-stained with a combination of lead and uranyl salts to add electron contrast before imaging in the SEM. ROIs selected in the fluorescence images were manually relocated in EM. The 2D LM image of the cell of interest (Fig.2e) was overlaid onto the 2D EM image (Fig.2f), enabling unequivocal identification of the nucleus, mitochondria and lysosomes in the EM images by the position of the fluorescent label. Though the nucleus and mitochondria are usually easy to identify in EM images by morphology alone, lysosomes are not, as they tend to have a heterogeneous appearance and content, within and between cell types. In this cell, both large (white boxes) and small (arrows) lysosomes could be unequivocally identified (Fig.2f and inset). MitoGFP was also successfully preserved in hiPSC-derived cortical neurons (Fig.2g) with electron contrast revealing mitochondrial networks across multiple cells (Fig.2h). Using the easyIRF protocol, the mitoplasm appeared dense and the cristae appeared light, meaning contrast was reversed compared to protocols incorporating osmium as a stain. Comparison of a magnified subregion shows matching mitochondrial patterns in both light and electron images from the same region.

Following successful proof-of-principle experiments using benchmark samples and probes, the distribution of ɑ-Syn protein was imaged 24 h after incubating monomers with hiPSC-derived cortical neurons (Fig.3). AF488-ɑ-Syn was preserved and visible in 200 nm sections from easyIRF blocks (Fig.3a). The 2D LM image was overlaid onto the 2D EM image (Fig.3b), revealing the distribution of ɑ-Syn throughout two cells (Fig.3c). ɑ-Syn was localised to several different cellular compartments, including mitochondria (as previously seen (Choi et al., 2022)) (Fig.2d, black arrows). The improved resolution and correlation precision of easyIRF sections also showed ɑ-Syn in close apposition to tortuous foldings of the nuclear envelope (Fig.2d, white arrows). A high density of ɑ-Syn was visible in large vacuolar structures with electron dense content (Fig.2e) and in smaller lysosome-like structures (Fig.2f) similar to the lysotracker-labelled structures seen in easyIRF images of HeLa cells (Fig.1f). In addition, small ɑ-Syn puncta were visible in the nucleus and cytoplasm. Despite the compromise in membrane preservation caused by the lack of osmium in the PLT step, the VP-CLEM-Kit workflow delivered vCLEM images that revealed insights into the subcellular localisation of pathogenic ɑ-Syn that could not be acquired through light microscopy alone.

To demonstrate that the easyIRF protocol also worked for larger biological samples, a kidney biopsy from a patient with immune-mediated transplant rejection was imaged (Fig.4). The biopsy was fixed, cut into 60 µm-thick slices using a vibratome, stained with Hoechst to highlight cell nuclei and labelled for CD34, a transmembrane phosphoglycoprotein expressed on endothelial cells, to highlight glomerular and peritubular capillaries. The tissue section was processed using a modified easyIRF protocol with extended times to accommodate the larger sample. The easyIRF block was then cut into 200 nm-thick sections, and the sections imaged using a commercial fluorescence microscope (Zeiss LSM900 Airyscan). Hoechst (Fig.4a) and AF568 (Fig.4b) were successfully preserved in resin. Following fluorescence microscopy, sections were post-stained and imaged in an electron microscope (Thermo Fisher Scientific Quanta 250 FEG SEM; Fig.4c). The fluorescence image was overlaid onto the EM (Fig.4d), with AF568-CD34 clearly highlighting the location of the endothelial cells of peritubular capillaries (PTCs) at low magnification across a large FOV. Ultrastructure and electron contrast was improved in the kidney tissue compared to the cell monolayer, possibly because the larger sample was less affected by solvent extraction during the PLT step. Features including rough endoplasmic reticulum (RER), mitochondria (M), collagen fibrils (CF), basement membrane (BM), cell junctions (CJ), tubular epithelial cells (TECs) and PTCs were clearly visible in the EM images (Fig.4e,f). The VP-CLEM-Kit thus adds significant ultrastructural context to fluorescence microscopy, as well as providing fast targeting to structures of interest in large tissue volumes for electron microscopy. This would enable, for example, a comprehensive assessment of the state of kidney transplant microvasculature to be built from the combination of peritubular capillary shape and density metrics derived from fluorescence with ultrastructural endothelial and basement membrane features.

The easyIRF protocol thus preserves a range of fluorescent proteins and dyes in different cell and tissue types. The overlay of mitoGFP onto mitochondrial structures as a benchmark gave confidence that fluorescence preservation was good enough to identify individual instances of labelled organelles in EM. Imaging the fluorescence and electron signals from the same ultrathin sections gave excellent registration of the two imaging modalities over wide fields of view, which is a significant improvement over pre-embedding vCLEM. This improvement was illustrated by the identification of a number of different organelle types containing pathogenic ɑ-Syn with localisation precisions below 1 micron, and the ability to quickly locate very thin endothelial cells within a large tissue section for targeted analysis of patient tissue.

### Serial sectioning for serial imaging in array tomography format

The strategy for moving easyIRF from two dimensions into three dimensions was to cut 200 nm-thick serial sections from the easyIRF blocks and to pick them up onto a conductive ITO-coated coverslip. The sections were then imaged sequentially to generate a series of images through the volume of the sample, first in the light microscope and then in the electron microscope. This process of sequential imaging of serial sections is called array tomography.

### Design and build of the tomoSTORM light microscope

At the time of building the VP-CLEM-Kit, there was no commercial light microscope designed to image ultrathin sections in array tomography format using widefield and super-resolution modalities. To deliver this new capability, a cost-effective user-friendly fluorescence microscope was designed and built, named tomoSTORM (Fig.5). This utilised the cost-effective modular *openFrame* microscope stand, complemented by low cost components including multimode diode lasers for *easySTORM* excitation (Kwakwa et al., 2016) and cooled CMOS cameras. The *openFrame* microscope stand was based on a stack of modular layers centred on the optical axis of the objective lens. The layers were locked together using dovetail connections and locking screws enabling them to be readily assembled, maintained and modified using standard tools. For tomoSTORM imaging, a motorised xy stage facilitated array scanning and tiling in conjunction with a motorised z-stage and *openAF* optical autofocus unit (Lightley et al., 2023). Reflected light imaging was implemented to aid in finding the ultrathin easyIRF sections.

**Figure 5.**
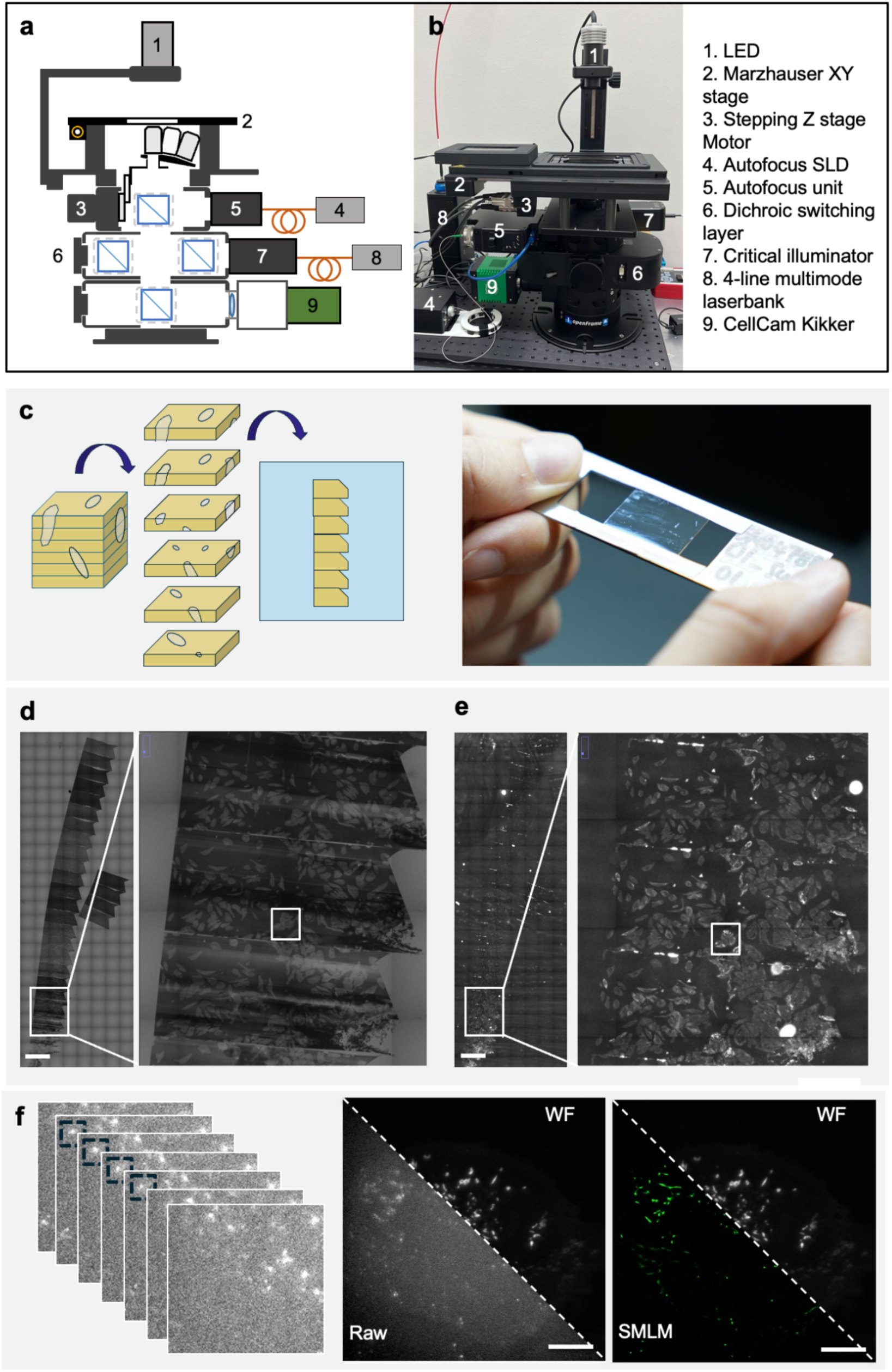
Sample mounting and light microscopy on the tomoSTORM microscope. (a) System diagram of the tomoSTORM open-hardware microscope based on the *openFrame* modular system, showing the hardware components used to acquire RF, WF and SMLM array tomography images. The system can switch between imaging modalities by manually switching the reflection and multi-line fluorescence dichroic cubes using the dichroic switcher layer. There is a manual 4-objective turret with 10x (Amscope, 10x/0.3 NA) and 100x (Amscope, 100x,/1.3 NA) objectives. The TEC-cooled CMOS camera is a CellCam Kikker (Cairn Research Ltd, CellCam Kikker). The system uses a XY stage (Märzhäuser Wetzlar GmbH, SCAN IM 130 x 85) and a Z-stepper motor (Cairn Research Ltd, Z-Act). The *openFrame* has an openAF autofocus unit to maintain focus during SMLM acquisitions. The excitation source is a 4-line multimode laserbank (Cairn Research Ltd, Laserbank) which is fibre coupled into the critical illuminator. (b) A photo showing the tomoSTORM light microscope with the individual hardware components labelled. (c) The easyIRF resin block was sectioned and section ribbons retrieved onto an ITO-coated coverslip. The coverslip was attached to a slide-sized metal mount and placed into the slide-holding insert on the xy stage for imaging. (d) RF and (e) WF tiled images were acquired with the 10x objective from a ribbon of easyIRF sections containing Hela cells expressing mitoGFP. Insets (white boxes) show sections 4-6 of the ribbon. Tiled WF images were recorded using the 100x/ 1.3 NA objective from section 4. (f) SMLM of mitoGFP in HeLa cells in an easyIRF section. Left: Multiple frames of a single FOV show blinking in sequential frames. Middle: A single frame of the raw microscope acquisition alongside WF, compared to Right: the final SMLM reconstruction in section 6 of the same sample, highlighting the resolution improvement achieved with SMLM.

easySTORM imaging (Kwakwa et al., 2016) on the tomoSTORM microscope provided SMLM acquisitions over large (>120 µm diameter) FOVs. The excitation was provided by a multiline laserbank comprising four multimode high power diode lasers (at 405 nm, 465 nm, 520 nm and 635 nm) that were coupled into the same large core (1000 µm) optical fibre. The optical fibre output was butt-coupled to a square core (1000 x 1000 µm) optical fibre that was imaged onto the sample plane to provide uniform critical illumination across the full FOV. An adjustable square aperture was placed in a conjugate image plane in the excitation path to enable the size of the FOV at the sample to be adjusted and to realise sharp edges of the FOV for tiled SMLM acquisitions. Low cost 10x and 100x objective lenses were used to image the sample at low and high magnification respectively. The imaging modality could be switched between RF (to locate the sections) and WF (to image fluorescent ROIs) modes using a manual dichroic beamsplitter switcher. A manual objective lens turret was used to switch between objective lenses. The sample was imaged using a cost effective thermoelectrically cooled CMOS camera (Cairn Research Ltd, CellCam Kikker), which has a pixel size of 4.63 µm. Image acquisition was automatically controlled using the multi-dimensional acquisition plug-in in Micro-Manager. Focus was maintained throughout the SMLM acquisitions and for slide scanning across the coverslip using the *openAF* hardware autofocus operating in single shot format to provide closed loop continuous focus correction during SMLM acquisitions.

### From 2D WF to 3D super resolution light microscopy with tomoSTORM

tomoSTORM was implemented to image ultrathin sections in RF mode, WF mode, and SMLM mode (Fig.5a,b). The ITO-coated coverslip holding the easyIRF sections was attached to a slide holder using tape (Fig.5c) and loaded onto the microscope stage. The motorised stage and software were designed to support tiling, so that a large FOV containing the section ribbons could be imaged with a 10x/ 0.3 NA objective lens. The rectangular aperture inside the critical illuminator unit prevented photobleaching of the fluorophores in areas outside the FOV, allowing accurate tessellation for tiled imaging. The tiled RF image revealed the section positions because the section edges were visible. The RF image also revealed the positions of cells within the sections due to the refractive index difference between the cells and the resin (Fig.5d). A tiled WF image was then acquired with the 10x/ 0.3 NA objective lens, revealing the distribution of fluorescence signals within cells in each section (Fig.5e), allowing the user to select ROIs for higher resolution imaging.

The tiled RF and WF images were montaged and then overlaid in Fiji to create an RF/WF map to support downstream navigation of sections and ROIs in the tomoSTORM microscope and EM. The 100x/ 1.3 NA oil objective lens was then used to image one section in WF mode to help select ROIs for subsequent imaging of the whole cell volume in SMLM mode across sequential sections, with a WF image of each ROI being recorded prior to each SMLM acquisition (Fig.5f).

Since fluorescent proteins and dyes blink in-resin, SMLM could be realised following the o*penFrame*-based implementation of *easySTORM*. To optimize blinking conditions for SMLM in resin-embedded mitoGFP and AF488, two excitation power titration experiments were conducted. For mitoGFP (Supp.Fig.1), an optimal power density of 330 W/cm² was identified as the best balance between signal intensity and background suppression. This power density aligned with previously reported data (Peddie et al.) and provided a reasonable fluorophore blinking rate while effectively driving most background fluorophores into the dark state. A similar power titration was performed for AF488 (Supp.Fig.2), yielding an optimal power density of 1231 W/cm². SMLM acquisitions were performed for each ROI (Fig.5f), and once complete, the stage was moved to image either a separate ROI within the same section or to the same ROI in the following section to build up a complete 3D SMLM volume from whole cells. SMLM acquisitions were reconstructed using Picasso.

### Tracking sections and fluorescent ROIs from LM to EM

The overlaid RF/WF image of the section ribbons from tomoSTORM was used to record section positions and select fluorescent ROIs for downstream imaging in EM (Fig.6). To support this process, a plugin was created in napari [https://doi.org/10.5281/zenodo.14719463] named napari microscopy array section setup (napari-mass) (Fig.6a). The project was first initialised (Fig.6b), followed by import of the overlaid RF/WF image. The layers were initialised (Fig.6c) to display the image data in the napari viewer (Fig.6d). The user could then draw around one of the sections in the viewer using the napari drawing tools. The section annotation was propagated to each sequential section using the copy and paste function (Fig.6e). The populate function was then used to extract each section and arrange them in a stack in the template view (Fig.6f). Using the template view, the fluorescence signal in each section was used as a guide to add ROI boxes around each cell of interest, which were then propagated through all sections. The scroll bar allowed the user to check that each selected ROI fully enclosed the cell of interest through the section stack (Fig.6g). Following section annotation and ROI selection, the positional data for both was exported as a json file. All plug-in functions were integrated into the user-friendly napari interface (Fig.6h). Propagation and templating concepts were inspired by the ‘propagation’ and ‘local view’ concepts (respectively) of the MagFinder Fiji plugin (https://github.com/templiert/MagFinder).

**Figure 6.**
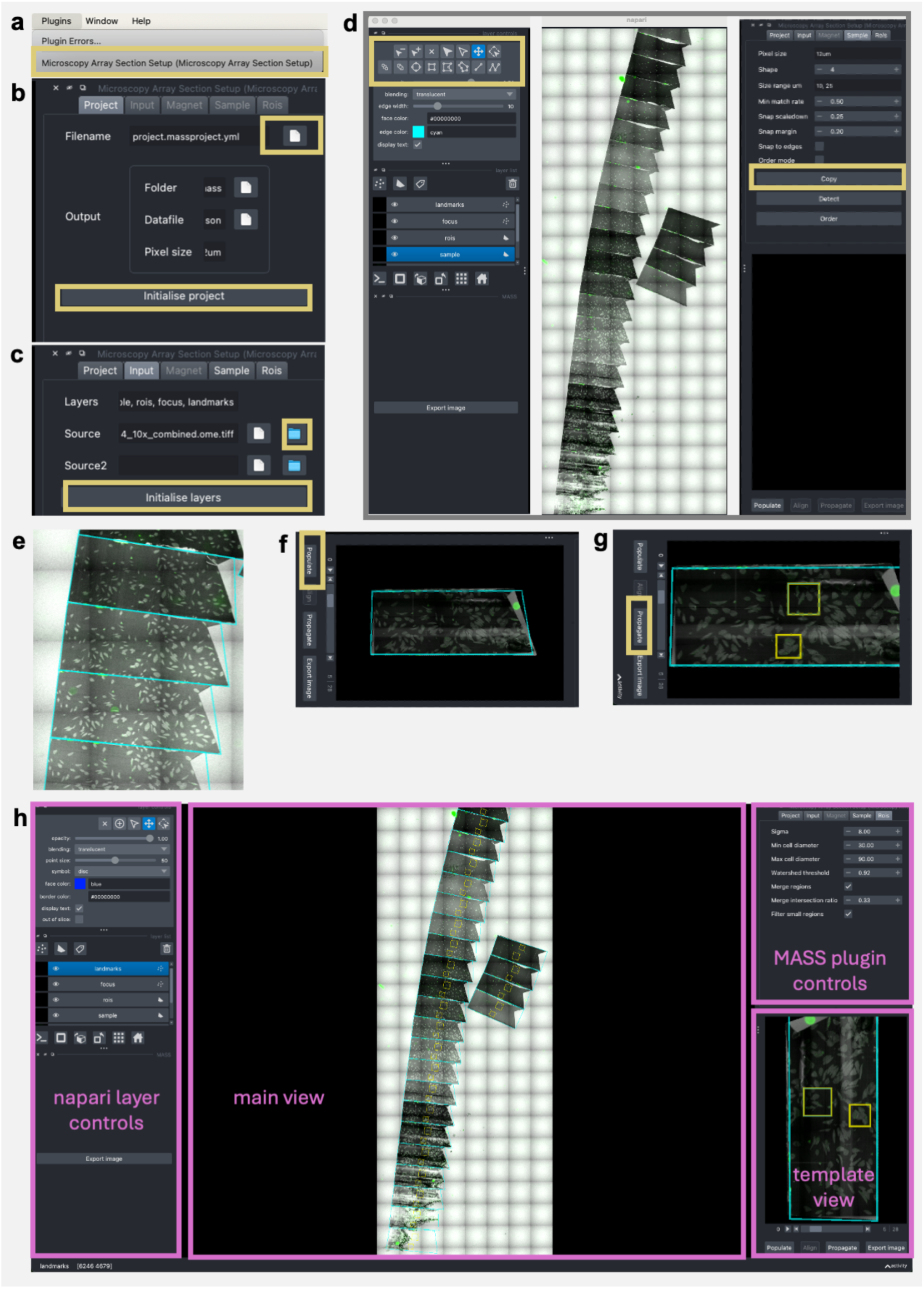
Section annotation and fluorescent ROI selection using the napari-mass plugin. (a) The microscopy array section setup (napari-mass) plugin is selected, allowing for (b) a project to be created and (c) layers initialised. (d) The RF and WF images acquired in the tomoSTORM light microscope are loaded as the source images and initialised. (e) A section is annotated using the napari drawing tools and the copy/paste controls used to propagate the annotation across all sessions in the image. (f) The section annotations are extracted and displayed as a stack in the template window, thus enabling (g) definition of fluorescent ROIs that are propagated axially through the serial sections. The annotations are stored in a format readable by SBEMimage for targeted imaging in the EM. (h) The workflow is implemented in the open-source napari platform.

### Post staining easyIRF sections for EM

Sections were post-stained for imaging in the Phenom Pharos tabletop SEM. Iterative improvements to the staining protocol resulted in the addition of 0.5% potassium permanganate to the uranyl and lead stains, which were subsequently used for all samples, whether imaging in the floor standing or tabletop SEMs.

### From 2D to 3D EM using array tomography

Array tomography is an established volume EM technique that generates a series of electron images from serial sections collected on a conductive substrate such as ITO-coated glass or silicon wafers. Array tomography is suitable for deployment in lower resource settings, since it can be performed using a field emission gun scanning electron microscope (FEG SEM) without significant modifications (White and Burden, 2023). However, even standard FEG SEMs are costly and require specialist environments to protect against acoustic and physical vibrations, temperature and airflow changes, and electromagnetic fields. To support wider access to the VP-CLEM-Kit, we instead used a significantly lower cost tabletop FEG SEM (the Thermo Fisher Scientific Phenom Pharos), which is user-friendly with a small footprint that can be positioned on a normal (albeit sturdy) table in a standard research laboratory environment (Fig.7). The system uses standard electrical connections that can be connected to an uninterruptible power supply for use in locations at risk of power outages. Sample loading is straightforward, with a short vacuum pumping cycle that takes two minutes due to the small size of the microscope chamber (Fig.7a).

**Figure 7.**
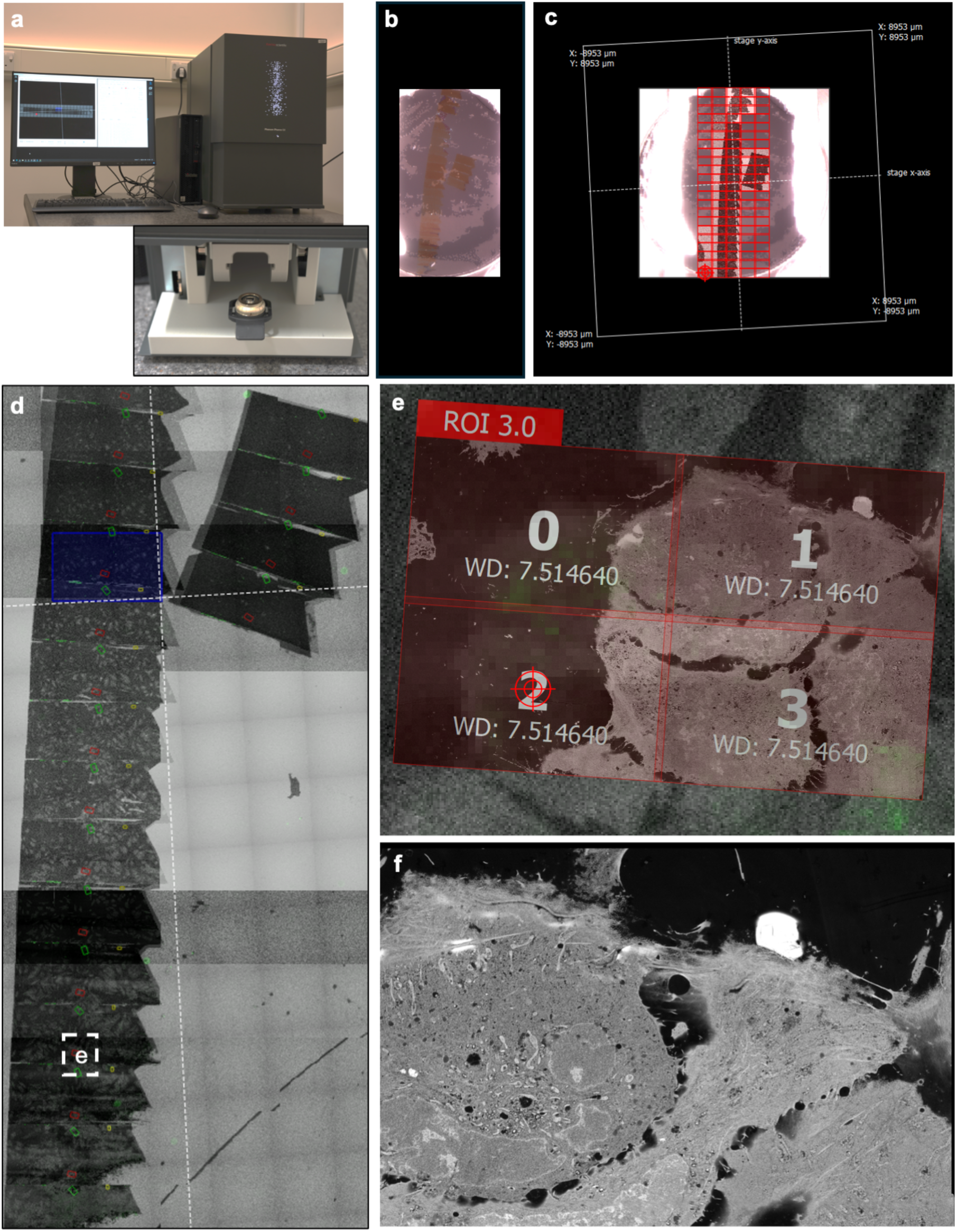
Array tomography on the Phenom Pharos tabletop SEM controlled by open-source SBEMimage software. (a) The Phenom Pharos is a cost-effective tabletop FEG SEM with a simple sample loading interface and user-friendly operation. (b) A stub overview image was acquired using the NavCAM, allowing detection of the sections. (c) A new array functionality was created in the open-source SBEMimage software, which controlled the Pharos through the API. The sections were selected in the NavCAM image by placing an imaging grid, the microscope was switched to electron imaging mode, and electron images of the sections were acquired. (d) The overlaid RF/WF image overlays with associated section and ROI annotations defined in the napari-mass plugin were imported into SBEMimage and aligned with the stub overview NavCAM and SEM images. (e) The ROIs defined in napari-mass were added as grids in SBEMimage using the ‘Create ROI’ tab, and (f) electron images were acquired from ROIs across sequential sections.

Manual operation of an array tomography workflow in a FEG SEM can be challenging, so an array tomography module was designed and integrated into the open-source SBEMimage software to simplify and control the process. The module builds and improves upon the existing MagC module of SBEMimage (Collectome LLC, Switzerland).

A stub overview image was acquired at 42x magnification using the in-microscope optical camera (NavCAM) to locate the serial sections (Fig.7b). The NavCAM image was used to define the area of the stub containing the sections (grid 0) in SBEMimage. An overview image of the sections was then acquired with the back scattered electron (BSE) detector in SBEMimage (Fig.7c). The overlaid RF/WF images and the json file containing xy coordinates of ROIs generated in napari-MASS were imported into SBEMimage. The overlaid RF/WF images were aligned to the NavCAM and EM stub overview images (Fig.7d). The list of ROI coordinates was then imported into SBEMimage and translated into the SEM space to generate grids to direct electron imaging across sequential sections (Fig.7e). A tiled image of each ROI was acquired from sequential sections at medium (20 nm pixels) and high (10 nm pixels) resolution (Fig.7f).

### EM image registration

Manual lateral alignment of EM image tiles was necessary where inaccuracies in stage movement or variations in focus and contrast caused tile mismatches. In these cases, the Fiji plug-in TrakEM2 was used for alignment. Tiles were imported and arranged within a single TrakEM2 layer, and positions were manually refined to ensure accurate alignment. Following the lateral stitching of image tiles within a section, TrakEM2 was used for axial alignment of the stack of tiled images. Each image was imported into a separate layer and manually adjusted by rotation and translation in the xy plane, ensuring alignment with the preceding section. For volume imaging datasets that were not significantly affected by image acquisition artefacts, a new pipeline for automated registration of image tiles was developed and applied, named muvis-align.

### Overlay of LM and EM images

To achieve precise overlay between LM (WF or SMLM) and EM images, a workflow using the BigWarp plugin in Fiji was developed. First, the corresponding LM and EM sections were imported into BigWarp, with the LM image designated as the ‘moving image’ and the EM image as the ‘target’. Typically, four or five well-distributed landmarks were selected to ensure an accurate transformation while minimizing artifacts. Once the transformation was applied, the aligned LM and EM sections were merged using Fiji’s merge channels function. This process was systematically repeated for each section. Since the EM images had already been axially aligned, the alignment of the LM sections naturally followed, ensuring consistency throughout the reconstructed volume.

### Increasing the resolution and volume of CLEM with SR-vCLEM

The full VP-CLEM-Kit pipeline was applied to three cells systems - HeLa cells expressing mitoGFP (Fig.8), hiPSC-derived cortical neurons expressing mitoGFP (Fig.9), and hiPSC-derived cortical neurons exposed to ɑ-Syn-AF488 (Fig.10). In each case, the fluorescent probes were preserved using the easyIRF protocol. Serial sections were cut and imaged using RF and WF modes with the 10x objective lens and WF mode with the 100x objective lens on the tomoSTORM microscope, before switching to SMLM mode. A 3D projection of the SMLM data was generated and colour-coded according to depth for each cell system (Fig.8a, Fig.9a, Fig.10a). Three representative planes are shown for each cell system to highlight resolution in the individual images (Fig.8b, Fig.9b, Fig.10b). For each cell, an individual section was then chosen to show a comparison of the WF (Fig.8c, Fig.9c, Fig.10c) and the SMLM (Fig.8d, Fig.9d, Fig.10d). Finally, EM images were acquired through the full cells (Fig.8e, Fig.9e, Fig.10e). Overlays of WF onto EM (Fig.8f, Fig.9f, Fig.10f) and SMLM onto EM (Fig.8g, Fig.9g, Fig.10g) allow visual assessment of the improvement in precision using SR-vCLEM.

**Figure 8.**
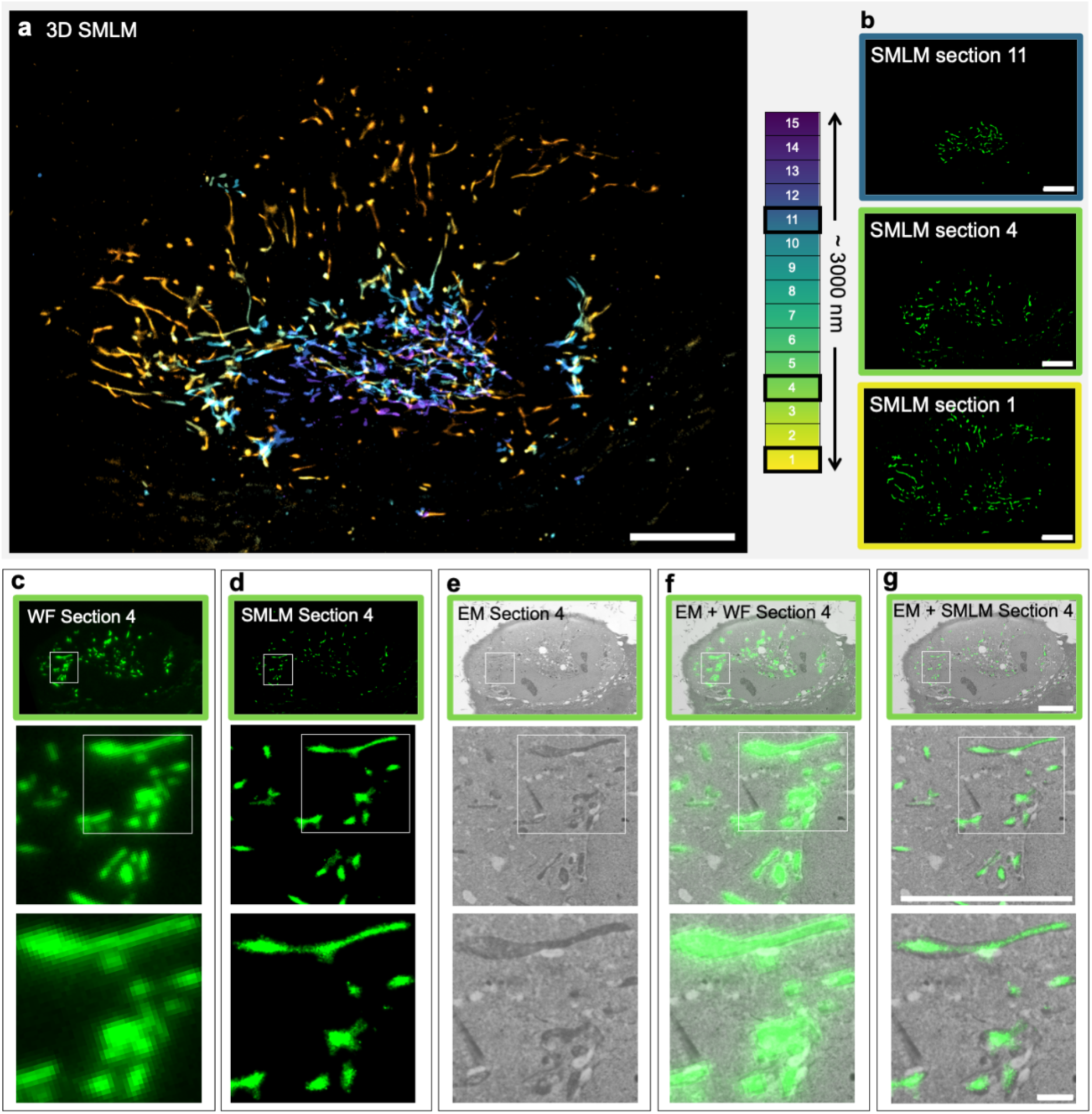
SR-vCLEM analysis of HeLa cells expressing mitoGFP, comparing WF-EM and SMLM-EM overlays for multimodal visualization of mitochondria. (a) 3D reconstruction of SMLM images generated from SMLM datasets acquired from 200 nm thick easyIRF sections through a total cell depth of 3000 nm. Each section was reconstructed using 10,000 frames. The 3D projection shows the spatial organization of mitochondria within the whole cell volume, showing continuity between planes. (b) Single-plane SMLM images corresponding to sections 1, 4, and 11, enabling depth-dependent visualization of mitochondrial structures with nanometric-scale resolution. (c) WF image of section 4, illustrating the overall mitochondrial distribution with diffraction-limited resolution. The estimated resolution, determined via decorrelation analysis, is 414 ± 13 nm, providing a broad overview of the labelled structures but lacking morphological details. (d) SMLM image of section 4, demonstrating a significantly improved spatial resolution compared to the WF image. The estimated resolution, determined via decorrelation analysis, is 84 ± 7 nm, enabling enhanced structural definition of mitochondrial networks. (e) SEM image of section 4. (f) Overlay of WF and EM images, facilitating correlation between mitochondrial localisation in the WF channel and ultrastructural features observed in EM. (g) Overlay of SMLM and EM images, providing improved spatial precision in the correlation of mitoGFP to mitochondrial ultrastructure. Scale bars represent 10 μm for all panels except for the panels on the bottom, where scale bars represent 1 μm.

**Figure 9.**
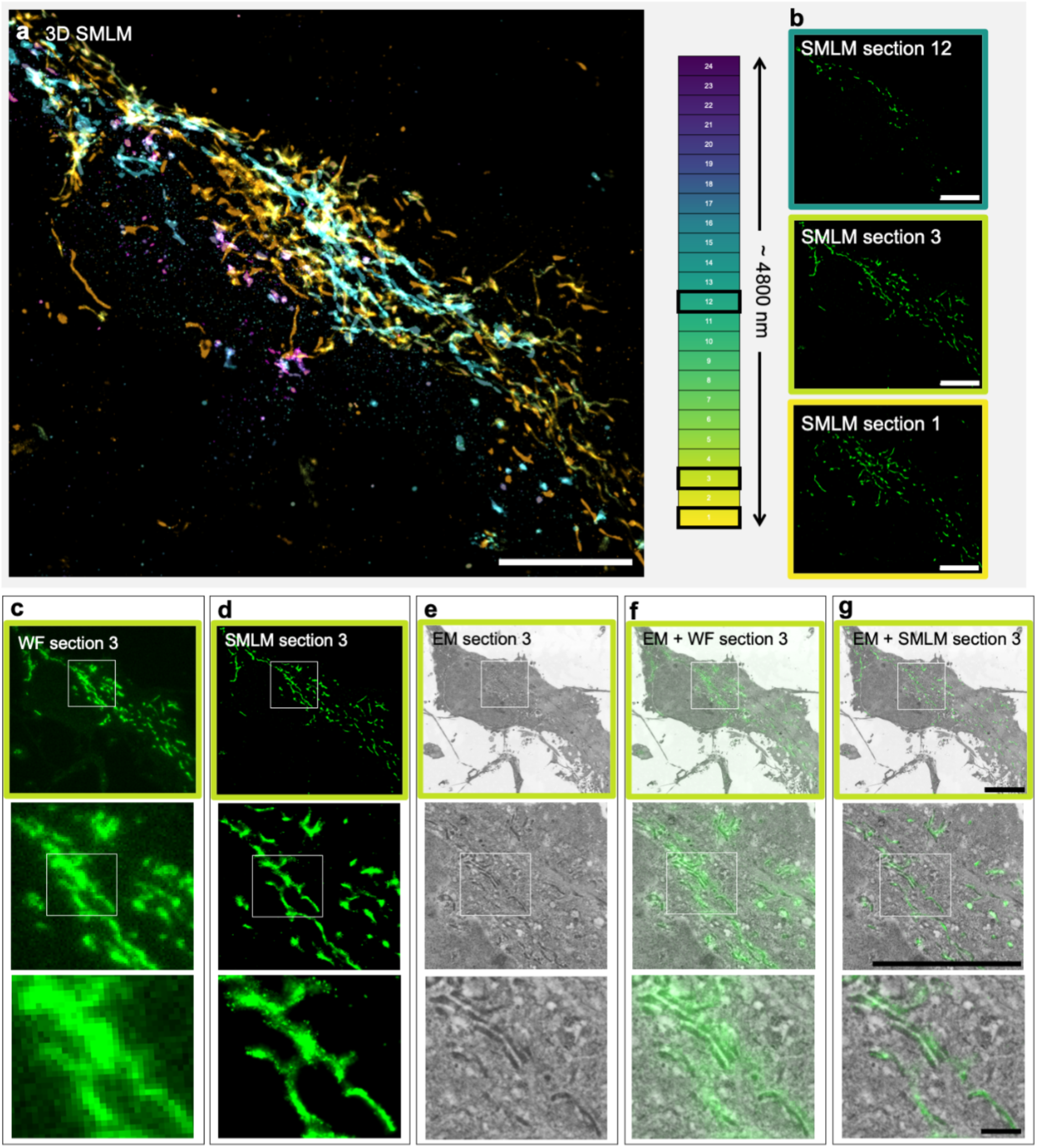
SR-vCLEM analysis of hiPSC-derived cortical neurons expressing mitoGFP, comparing WF-EM and SMLM-EM overlays for multimodal visualization of mitochondria. (a) 3D reconstruction of SMLM images generated from SMLM datasets acquired from 200 nm thick easyIRF sections through a total cell depth of 4800 nm. Each section was reconstructed using 10,000 frames. The 3D projection shows the spatial organization of mitochondria within the whole cell volume, showing continuity between planes. (b) Single-plane SMLM images corresponding to sections 1, 3, and 12, enabling depth-dependent visualization of mitochondrial structures with nanometric-scale resolution. (c) WF image of section 3, illustrating the overall mitochondrial distribution with diffraction-limited resolution. The estimated resolution, determined via decorrelation analysis, is 688 ± 75 nm, providing a broad overview of the labelled structures but lacking morphological details. (d) SMLM image of section 3, demonstrating a significantly improved spatial resolution compared to the WF image. The estimated resolution, determined via decorrelation analysis, is 78 ± 7 nm, enabling enhanced structural definition of mitochondrial networks. (e) SEM image of section 3. (f) Overlay of WF and EM images, facilitating correlation between mitochondrial localisation in the WF channel and ultrastructural features observed in EM. (g) Overlay of SMLM and EM images, providing improved spatial precision in the correlation of mitoGFP to mitochondrial ultrastructure. Scale bars represent 10 μm for all panels except for the panels on the bottom, where scale bars represent 1 μm.

**Figure 10.**
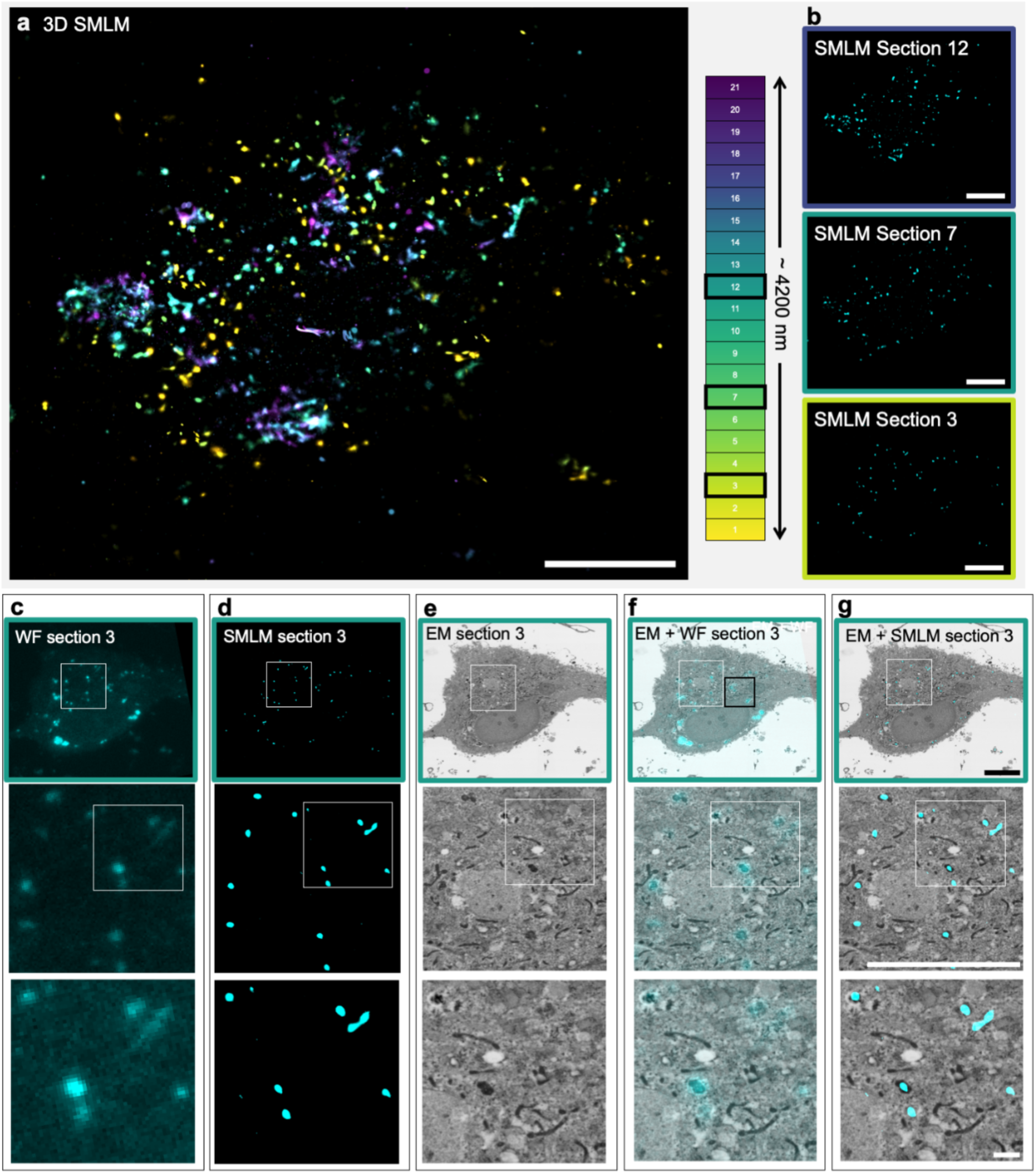
SR-vCLEM analysis of hiPSC-derived cortical neurons exposed to ɑ-Syn-AF488, comparing WF-EM and SMLM-EM overlays for multimodal visualization of ɑ-Syn localisation. (a) 3D reconstruction of SMLM images generated from SMLM datasets acquired from 200 nm thick easyIRF sections through a total cell depth of 4200 nm. Each section was reconstructed using 10,000 frames. The 3D projection shows the spatial organization of ɑ-Syn within the whole cell volume, showing continuity between planes. (b) Single-plane SMLM images corresponding to sections 3, 7, and 12, enabling depth-dependent visualization of ɑ-Syn with nanometric-scale resolution. (c) WF image of section 3, illustrating the overall ɑ-Syn distribution with diffraction-limited resolution. The estimated resolution, determined via decorrelation analysis, is 783 ± 72 nm, providing a broad overview of the labelled structures but lacking morphological details. (d) SMLM image of section 3, demonstrating a significantly improved spatial resolution compared to the WF image. The estimated resolution, determined via decorrelation analysis, is 70 ± 8 nm, enabling enhanced structural definition of ɑ-Syn distribution. (e) SEM image of section 3. (f) Overlay of WF and EM images, facilitating correlation between ɑ-Syn localisation in the WF image and ultrastructural features observed in EM. (g) Overlay of SMLM and EM images, providing improved spatial precision in the correlation of ɑ-Syn to cellular ultrastructure.

As expected, mitoGFP localised to mitochondria in WF/EM and SMLM/EM overlays from HeLa cells and iPSC-derived cortical neurons. These benchmark experiments confirmed that SR-vCLEM effectively localised fluorescently-tagged proteins to organelles (Fig.8f,g; Fig.9f,g). To quantify the accuracy of FM-to-EM alignment, we measured the overlay precision using the Jaccard index (JI), a metric that assesses the similarity between two sets by calculating the ratio of their intersection to their union. JI was computed across all sections of each sample by comparing a manually annotated EM mitochondrial mask against thresholded SMLM and WF images. The final reported values represent the mean and standard deviation of JI measurements across all sections.

In HeLa cells expressing mitoGFP, the overlay improved significantly from JI = 0.26 ± 0.06 in WF mode to JI = 0.40 ± 0.09 in SMLM mode (Fig.8), demonstrating a notable enhancement in registration accuracy. In hiPSC-derived cortical neurons expressing mitoGFP (Fig.9), the improvement was more modest, increasing from JI = 0.32 ± 0.03 (WF) to JI = 0.39 ± 0.04 (SMLM). The relatively low Jaccard index values across both modalities reflect inherent challenges in precisely annotating mitochondria within resin-embedded samples, where contrast limitations and morphological variability complicate segmentation. Nonetheless, all visually identifiable mitochondria exhibited corresponding SMLM signals. Notably, the SMLM overlay also revealed additional mitochondrial structures that were less apparent in WF imaging, confirming the method’s ability to enhance structural identification and provide valuable insights.

The comparison between SMLM and WF in HeLa cells expressing mitoGFP (Fig.8) revealed a strong structural correlation, with a resolution-scaled Pearson coefficient (RSP) of 0.70 ± 0.08, as computed using SQUIRREL (Culley et al., 2018), further validating the reliability of the approach. The RSP measures the correlation between super-resolved and diffraction-limited images while accounting for differences in resolution, giving an indication of reconstruction fidelity. The SMLM reconstruction achieved a resolution estimation through decorrelation analysis (Descloux et al., 2019) of 84 ± 14 nm, representing a substantial improvement over the corresponding widefield estimation of 371 ± 38 nm. Note that this decorrelation measurement provides an estimation of the scale of the smallest structures present within the image, rather than the best spatial resolution achievable with any sample. A similar enhancement was observed in hiPSC-derived cortical neurons expressing mitoGFP (Fig.9), where SMLM reconstruction achieved a resolution estimate of 78 ± 14 nm, compared to the widefield resolution estimate of 492 ± 67 nm, while maintaining an RSP of 0.63 ± 0.18, demonstrating the consistency of the reconstruction.

For ɑ-Syn-AF488 (Fig.10), the signal localized to both large and small lysosome-like structures, with overlay precision significantly improving from JI = 0.20 ± 0.10 in WF mode to JI = 0.41 ± 0.07 in SMLM mode. The SMLM reconstruction achieved a resolution estimate of 70 ± 16 nm, significantly surpassing the corresponding widefield resolution estimate of 582 ± 95 nm. Due to the complexity of the sample, the RSP was lower (0.58 ± 0.07), likely due to reduced blinking efficiency or elevated background in certain sections. With the improved correlation precision delivered by SR-vCLEM, it was possible to localise ɑ-Syn inside the lumen of small lysosome-like structures (Fig.10g), which was a significant improvement over the WF localisation (Fig.3d), where the overlay of fluorescence onto the electron image showed ɑ-Syn localisation inside and outside the lysosome-like structures due to the lower resolution of the WF image. Future SR-vCLEM experiments combining organelle tracker dyes (e.g. lysotracker, mitotracker) with ɑ-Syn-AF488 in the same sample should allow further dissection of ɑ-Syn processing in different subcellular locations.

All reported values were computed across sections, with statistical values derived from the full dataset. Improvements in resolution in all samples highlight the strength of SR-vCLEM in refining the precision of correlation of functional labels to underlying subcellular structure. In terms of cell biology, the improvement means that molecules can be localised to organelles smaller than one micron (e.g. early endosomes, trafficking vesicles, centrioles, viruses) and subdomains of larger organelles, as well as providing the possibility to differentiate between lumenal and cytosolic localisations for membrane-proximal molecules (demonstrated here by ɑ-Syn localisations at the cytosolic face of the nuclear envelope). This capability will enable important questions to be addressed in disease biology.

In summary, the VP-CLEM-Kit pipeline delivered approximately 5-fold enhanced resolution in LM by leveraging the blinking of fluorophores and SMLM, with a robust RSP (indicating high-quality super-resolution reconstruction) and significantly improved overlay accuracy compared to WF images (measured by the JI). As well as new capability in SR-vCLEM, the VP-CLEM-Kit also improved usability and affordability, costing 7-fold less than the existing HPF IRF workflow.

## Discussion

As all forms of microscopy move from two to three dimensions, and imaging across scales connects large fields of view with high resolution detail, correlative workflows are becoming more common. However, the majority of correlative workflows that reach ultrastructural resolution are performed by EM experts rather than LM experts. Since CLEM uses both imaging modalities, one would aspire to the technique being accessible to both light and electron microscopy facilities to benefit both communities. EM adds the context of membrane and organelle structure to fluorescence-based molecular localisations for light microscopists, and LM adds identity and molecular distributions to cell structures for electron microscopists. There are a few reasons why CLEM has not been broadly adopted by light microscopists, including the expense and complexity of the instrumentation, as well as considerations of the toxicity of the sample preparation protocols required for electron imaging.

In this work, we present a new correlative workflow that removes many of the barriers to broader adoption of CLEM. The easyIRF protocol minimises the requirement for costly specialist sample preparation equipment and the more toxic chemicals, with lower cost light and electron microscopes then being able to provide SR-vCLEM capability, thereby widening access to visual proteomics research. Nevertheless, the pipeline currently still requires access to an ultramicrotome for cutting ultrathin sections.

To support light microscopy facilities and research labs to adopt the VP-CLEM-Kit, ultramicrotomy could be performed in collaboration with a local or remote EM facility, since the samples are stable once embedded in resin, and both blocks and sections can be easily shipped. In addition, recent advances in commercial ultramicrotomes, including partial automation of the alignment of the diamond knife to the block face, support accessibility of ultramicrotomy for new users. A number of solutions have been developed for automated section retrieval, some of which have now been commercialised (Fulton et al., 2024; Hayworth et al., 2006; Li et al., 2017; Templier, 2019). In future iterations of the VP-CLEM-Kit, it is likely that automation of ultramicrotomy will enable yet broader adoption of the technology by removing the need for an expert electron microscopist to be involved in the pipeline.

In this work, ultrathin sections were cut at 200 nm thickness. In future work, SR-vCLEM sections of less than 200 nm thickness could be explored. We expect that thinner sections would result in improved resolution in both fluorescence and electron images, but with a trade-off in contrast due to the lower number of fluorescent molecules and stained membranes present in the thinner sections. It seems likely that 50 nm isotropic resolution could be achievable using tomoSTORM on 50 nm sections. Such effective precision of localised labels would approach that of a primary and secondary antibody pair with a 10 nm gold label traditionally used in EM to localise proteins to structures. SR-vCLEM would then be a highly attractive alternative to EM immunolabelling, especially since there is a lack of EM-compatible antibodies for most cell and tissue types, except for the brain, where the connectomics community have made a concerted effort to screen and/ or develop reagents (Micheva et al., 2023). An additional consideration is that an increased number of thinner sections for a given volume will increase data acquisition times and generate more data to be processed and stored. We suggest that section thickness should therefore be chosen according to the research question being asked, and the properties of the sample (probe labelling density, size of target structure, volume to be imaged etc).

GFP family probes are some of the most commonly used in bioscience research, but GFP does not usually blink well enough for conventional SMLM in liquid buffers. Thus IRF and easyIRF allow researchers to achieve super resolution in GFP-labelled cells and tissues without having to re-clone and re-characterise cell lines and animal models. We note that GFP and Alexa-dyes blink under the same conditions in IRF and easyIRF sections and it should therefore be possible to image multiple probes in the same section using SMLM for colocalisation studies and for deeper visual proteomics.

The easyIRF protocol avoids the use of glutaraldehyde in the fixative, heavy metals in the dehydration solution, and epoxy resins for embedding. This lowers the toxicity of the protocol, and also minimises fluorophore quenching or masking, thus maximising the number of fluorescent molecules available for SMLM in the sections. As well as providing optimal in-resin detection of low-expressing fluorescent constructs, this means that the easyIRF protocol is likely to offer benefits even when using fluorescent probes that have been reported to have some resilience to glutaraldehyde, osmium and epoxy resins. For these probes, fluorescence emission is still reportedly reduced to between ∼10-60% of the original levels once osmicated and embedded in epoxy resin, making them unsuitable for in-resin SMLM (Peng et al., 2022; Sanada et al., 2022; Tanida et al., 2020b; Tanida et al., 2023)

The lack of glutaraldehyde and osmium in the easyIRF protocol results in the main compromise of the method - that of reduced visibility of membranes in the sample for electron imaging. It is unclear whether the lack of membrane visibility in the images is due to extraction of lipids during the PLT dehydration, or due to lower binding of heavy metal stains to the membranes, as this happens after the samples are embedded in resin. Efforts to address membrane contrast should be prioritised in future work, because once membranes are clearly visible, SR-vCLEM using the easyIRF protocol could become the main CLEM method due to superior performance in resolution and volume. Since the VP-CLEM-Kit is both accessible and cost effective, efforts to solve the ‘contrast compromise’ can progress more quickly because a larger community of imaging scientists will be able to take part.

Though membrane visibility was reduced using easyIRF, other components of the cell were well stained and clearly visible, including the nucleus, nuclear pores in the nuclear envelope, chromatin, nucleoli, cytoskeleton, and content in lysosomes. Mitochondria were clearly visible, though with reversed contrast, as the nucleoplasm appears dense and the cristae appear light. Lipid droplets, endoplasmic reticulum, endosomes and Golgi apparatus were more difficult to identify. As with all research techniques and technologies, the method used should be matched to the research question and the sample. Until membrane contrast is improved, easyIRF may be unsuitable for answering questions about membrane-membrane contact sites, while still being suitable for investigations of mitochondrial networks and lysosomal trafficking, for example. Though the mechanism is not yet understood, it is intriguing that tissue ultrastructure appears to be better preserved and contrasted than for cell monolayers, making SR-vCLEM relevant for a wide range of larger biological samples.

Though the VP-CLEM-Kit was designed for democratised SR-vCLEM, it can also deliver advantages when broken down into its constituent parts. Using the easyIRF protocol and the tomoSTORM light microscope alone could deliver close to 50 nm isotropic resolution through any volume, limited only by the number of sections cut and imaged. This volume LM technique would leverage the blinking of standard fluorescent proteins in cells and tissues when embedded in resin. To reach very large sample volumes at low cost, stripped-back versions of the tomoSTORM microscope could be built that have only the SMLM imaging mode, and these could be run in parallel. Similarly, using SBEMimage to control array tomography on the Phenom Pharos tabletop SEM alone could enable volume EM in most EM facilities, even in lower resource settings, using standard heavy metal stained, epoxy resin embedded samples. There is also scope in the future to design and build a customised tabletop FEG SEM with a larger chamber and further automation of array tomography acquisition software for larger volume, cost effective, user-friendly volume EM.

Challenges in data handling and analysis remain. Registration of serial electron images in x, y and z is slow when performed manually using TrakEM2, especially when handling big image data. Our new automated registration pipeline (muvis-align) can fail due to acquisition artefacts such as folds in sections and inaccurate stage movements during image tiling. Even though the light and electron images are acquired from the same section, the overlay of the images from the two imaging modalities becomes more challenging as the resolution of the light microscopy increases. We propose that inclusion of off-target landmark labels would support more accurate overlay of SMLM data onto EM data for protein localisation and organelle identification. As reported in our previous work, mitochondria are useful as cellular landmarks for alignment as they tend to be well-distributed throughout the cell (Krentzel et al., 2024). Further development of automation in the muvis-align pipeline and addition of a user-friendly interface would facilitate both registration and correlation for SR-vCLEM.

The VP-CLEM-Kit provides insights into cellular context that are not normally available to light microscopists. Imaging of biopsy tissue from human patients also suggests that the VP-CLEM-Kit may prove useful for both clinical research, and perhaps eventually to introduce CLEM into diagnostic settings due to its relatively low cost and ease of use. Perhaps most importantly, the VP-CLEM-Kit has the potential to engage both light and electron microscopists in the same technical space, thereby supporting growth in interdisciplinary imaging and analysis skillsets, and a scientific workforce better equipped to push the boundaries of imaging across scales.

## Materials and Methods

### HeLa cell culture, constructs and transfection

Human cervical cancer epithelial (HeLa) cells were obtained from the American Type Culture Collection (ATCC, CCL-2). Cells were maintained in Dulbecco’s Modified Eagle Medium (DMEM) supplemented with 10% Gibco Fetal Bovine Serum (FBS) (Thermo Fisher Scientific) and 1% penicillin/streptomycin (Pen/Strep) (Sigma Aldrich). For experiments, HeLa cells were seeded overnight onto two 10 mm glass coverslips per 3.5 cm-diameter plastic culture dish at a density of 150,000 cells per dish. Thereafter, cells were transfected with 0.5 µg of mitoGFP construct (pLV-mitoGFP # 44385, Addgene) per dish, using Lipofectamine LTX and PLUS reagent (Invitrogen, Life Technologies Ltd, Paisley) in Opti-MEM medium (Gibco, Life Technologies Ltd, Paisley) according to the manufacturer’s instructions. The transfection mix was added to the cells in antibiotic-free medium and incubated for 18–24 h. Thereafter, cells were either fixed or stained with 2 µg/µl of Hoechst 33342 and 150 nM of Lysotracker red (Molecular Probes, Thermo Fisher Scientific) for 30 mins in culture medium, washed 3 times with culture medium and then fixed.

### Human-induced pluripotent stem cell culture

hiPSCs were maintained at 37°C and 5% CO_2_ in feeder-free monolayers on Geltrex-coated (Thermo Fisher Scientific) cell culture plates and were fed daily with mTeSR 1 media (StemCell Technologies). For passage, hiPSCs were incubated with 0.5 mM EDTA (Life Technologies). hiPSCs underwent mycoplasma testing and short tandem repeat sequencing to confirm hiPSC line identity.

### Cortical neuron differentiation and treatment

hiPSC-derived cortical neurons were differentiated using a previously published protocol (Choi et al., 2022; Shi et al., 2012). Briefly, neural induction was initiated through the addition of dual SMAD inhibitors, SB431542 (10 µM; Tocris) and dorsomorphin dihydrochloride (1µM; Tocris), to N2B27 medium. N2B27 medium consisted of DMEM/F12, insulin, 2-mercaptoethanol, non-essential amino acids, Pen/Strep, Glutamax, N2 and B27 supplements. After 10 days of dual SMAD inhibition, neuroepithelial sheets were dissociated using dispase and the cells were then further cultured in N2B27 medium with twice weekly feeding until >70 days of differentiation. For final plating, cells were dissociated into a single-cell suspension using accutase, and were seeded onto two poly-D-lysine (PDL)- and laminin-coated coverslips per 3.5 cm-diameter plastic culture dish at a density of 80,000 cells/cm^2^. hiPSC-derived cortical neurons were then either transduced with mitoGFP lentivirus to fluorescently tag the mitochondria or treated with monomeric ɑ-Syn protein. ɑ-Syn was purified from *Escherichia coli and* fluorescently labelled with Alexa Fluor dyes via the cysteine thiol moiety as previously described (Hoyer et al., 2002; Thirunavukkuarasu et al., 2008). hiPSC-derived cortical neurons were fixed 24 hours after ɑ-Syn treatment. We have previously reported the aggregation kinetics of Alexa Fluor-labelled ɑ-Syn in hiPSC-derived cortical neurons (Choi et al., 2022).

### Human kidney biopsy collection and processing

Material excess to diagnostic needs from indication kidney transplant biopsy samples was taken at Imperial College NHS Trust Renal Unit. Imperial College Healthcare NHS Trust Tissue Bank has ethical approval to both collect human tissue excess to diagnostic needs and release material to researchers (MREC 22/WA/2836). Patients undergoing a procedure at Imperial College Healthcare NHS Trust were asked at time of the procedure if they consented to material surplus to diagnostic needs from their tissues being used for research, using PIS v.8. Their response was recorded in the electronic patient record and on the pathology request form. Approval for our specific project was registered with Imperial College Healthcare Tissue Bank as project R14094 “Outcome analysis after renal transplantation in patients with de novo donor-specific antibodies”.

Biopsy samples were fixed in 4% EM-grade formaldehyde (TAAB) in 0.1 M phosphate buffer (PB) pH 7.4 and stored at 4°C. Samples were transferred to 1% formaldehyde in 0.1 M PB and this storage fixative was replaced approximately every 3 months. For experiments, biopsies were embedded vertically in 3% low melting point agarose (Sigma Aldrich) in 0.1 M phosphate buffer (PB) and 60 µm transverse vibratome slices were cut using a VT1200 vibrating blade microtome (Leica Biosystems). Vibratome settings were 0.3 mm/s speed of advance and 0.7 mm amplitude, and the blade was carbon steel (FEATHER®). Slices were collected and transferred in sequence to an optical bottom 24-well µ-Plate (Ibidi) filled with 0.1 M PB. Vibratome slices were stored in 1% FA in 0.1 M PB at 4°C.

All CD34 immunofluorescence labelling and Hoechst staining steps were performed in sterile 24-well plates, at room temperature on a plate shaker, unless otherwise specified. Vibratome slices were washed 3 times in 0.1 M PB then incubated in 0.05 M glycine (Sigma Aldrich) in 0.1 M PB for 30 mins. Slices were washed for 30 mins in 0.1 M PB then incubated in a blocking solution of 1% bovine serum albumin (BSA; Sigma Aldrich) in 0.1 M PB for 30 mins. Slices were incubated in the primary antibody (mouse monoclonal [QBEnd 10] to CD34 [Agilent]) diluted 1:30 in 0.1 M PB overnight at 4°C without agitation. Slices were washed 3 times in 0.1 M PB then incubated in the secondary antibody (goat anti-mouse IgG H&L Alexa Fluor® 568 [abcam]) diluted 1:100 plus 4 µg/ml Hoechst 33342 (Invitrogen Molecular Probes) in 0.1 M PB for 60 mins. Slices were washed for 30 mins in 0.1 M PB.

### Sample processing of cell monolayers using the easyIRF protocol

HeLa cells and hiPSC-derived cortical neurons were fixed in double strength fix (8% methanol-free formaldehyde in 0.4 M HEPES, pH 7.4, pre-warmed to 37°C) added 1:1 (v:v) to the growth medium for 15 mins at room temperature. Cells were then fixed in single strength fix (4% methanol-free formaldehyde in 0.2 M HEPES) for 24 hours at 4°C and washed with 2 ml of 0.2 M HEPES, 3 x 5 min on ice. Coverslips were transferred to metal dishes containing precooled 50% acetone and transferred to the automated freeze substitution unit (AFS2; Leica Microsystems, Vienna) for dehydration, infiltration and polymerisation. Cells were dehydrated in 50% acetone while the temperature was lowered from 4°C to -20°C over 20 min. Cells were then further dehydrated in 70%, 90% and 2x anhydrous acetone for 10 min each step at -20°C. LR White resin (pre-catalysed LRW; Agar Scientific #AGR1280) was mixed with benzoin methyl ether (Sigma Aldrich) to a final concentration of 0.75%, and a graded series of resin in acetone prepared at 25%, 50% and 75% (v/v). Cells were infiltrated with the resin:acetone series for 10 mins for each step followed by 100% LR White overnight, all at -20°C in the dark. Cells were further infiltrated for 2h at -20°C in the dark in freshly prepared 100% LR White (pre-cooled at 4 and then at -20°C). A disc cut from an aclar sheet was placed on top of the aluminium dish to exclude oxygen, and the resin was polymerised under 360 nm UV light for 48 h before warming to 4°C over 16 hours. The aluminium dish was peeled away from the polymerised resin block, and a jeweller’s saw was used to cut around the coverslip. The block was partially submerged into liquid nitrogen with the coverslip facing downwards until the glass separated from the resin surface, leaving a monolayer of cells at the surface of the resin block. Blocks were stored in the dark at 4°C.

### Sample processing of kidney biopsies using a variation of the easyIRF protocol

After CD34 immunofluorescence staining, kidney slices were stored in 1% FA in 0.1 M PB at 4°C. For low temperature dehydration and resin-embedding, kidney slices were added to a metal dish containing 4°C 50% ethanol which was transferred to the automated freeze substitution unit (AFS2; Leica Microsystems). Kidney tissue was dehydrated in 50% ethanol while the temperature was lowered from 4°C to -20°C over 20 min. Kidney slices were kept at -20°C and further dehydrated in 70% ethanol for 30 min, 90% ethanol for 60 min and 2x anhydrous ethanol for 60 min each. LR White resin (pre-catalysed LRW; Agar Scientific #AGR1280) was mixed with benzoin methyl ether (Sigma Aldrich) to a final concentration of 0.75%, and a graded series of resin in ethanol prepared at 25%, 50% and 75% (v/v). Kidney slices were infiltrated with the resin:ethanol series for 30 mins for each step followed by 100% LR White overnight, all at -20°C in the dark. The AFS chamber temperature was increased to 4°C over 20 min and kidney slices were further infiltrated for 30 min at 4°C in the dark in freshly prepared 100% LR White. The AFS chamber temperature was decreased back to -20°C over 20 min. A disc cut from an aclar sheet was placed on top of the aluminium dish to exclude oxygen, and the resin was polymerised under 360 nm UV light for 48 h before warming to 4°C over 16 hours. The aluminium dish was peeled away from the polymerised resin block, and a razor blade was used to cut around the kidney slices, being careful not to crack the brittle resin. Blocks were stored in the dark at 4°C.

### Serial ultrathin sectioning

A piece of the polymerised block (∼2 mm^2^) was cut out using a razor blade and mounted onto a dummy resin block with superglue. In order to produce neat ribbons, the block was trimmed to produce a parallel leading and trailing edge, with asymmetric sides for orientation. A 90° diamond trimming knife (Diatome) was used to trim the sides of the mesa so that the section size stayed the same across the ribbon. Serial 200 nm-thick sections were cut using a UC7 ultramicrotome (Leica Microsystems, Vienna), collected onto a glow discharged ITO-coated glass coverslip (Delmic B.V., Delft) using the paper clip method (White and Burden, 2023). The coverslip was attached to a slide holder with tape for imaging with a light microscope.

### Imaging easyIRF sections using a commercial confocal light microscope

An AxioObserver 7 LSM900 with Airyscan 2 microscope with Zen 3.1 software (Carl Zeiss Ltd) was used to image easyIRF samples during the development phase of the easyIRF protocol. The ITO-coated coverslips with sections was attached to a slide holder using tape. Section ribbons were first imaged using a Plan-Apochromat 20x/0.8 NA objective lens with widefield reflection light using a tile scan to map ribbons followed by imaging the same region with fluorescence mode to check fluorescence preservation. A reflection filter cube was created using the Zeiss type 00 filter cube, where the excitation filter was removed and replaced with a neutral density filter to significantly reduce the illumination intensity, whilst keeping the LP 615 emission filter. Following low magnification imaging, one section was tiled and imaged at high resolution in AiryScan super-resolution (SR mode) mode using a Plan-Apochromat 63x/1.4 NA oil objective lens. Imaging parameters were set using the smart setup tool, and channels (Alexa dyes, Lysotracker, mitoGFP and Hoechst) were acquired sequentially to limit crosstalk. Airyscan images were processed using the AiryScan processing tool in Zen Blue (Carl Zeiss AG).

### Post-staining sections to add electron contrast

After imaging with light microscopy, the coverslip was removed from the slide holder and cleaned with ethanol to remove oil. The sections were baked at 60° for 10 mins on a hot plate to facilitate attachment (section-side-up). Sections were post-stained with 2% uranyl acetate in ddH_2_O (w/v) for 10 mins at room temperature in the dark. Coverslips were then floated on a droplet of water for 30 secs (section-side-down) followed by dipping the coverslip 10 times in each of five beakers of water. Sections were then stained in Reynolds lead citrate for 10 min and washed as before. Coverslips were air dried, mounted onto an SEM stub with a carbon sticky tab and the edges coated with silver paint on the sides to improve conductivity.

### Imaging easyIRF sections using SEM

Following post staining, the same section imaged with the Zeiss Airyscan was imaged on the FEI Quanta 250 FEG SEM (Thermo Fisher Scientific) or Zeiss Gemini 460 (Carl Zeiss). For imaging, both systems were used at high vacuum. On the Quanta 250 FEG SEM, FEI MAPS software was used to acquire tiled images. The vCD backscatter detector was used at a working distance of 5 mm (cells) or 6 mm (biopsy tissue), and inverted contrast images were acquired at an accelerating voltage of 3 kV, spot size 2.5, 30 µm aperture, and dwell time of 60 µs for a 3072 x 2048 image frames or a dwell time of 30 µs for a 1536 x 1024 image frames for cells, or an accelerating voltage of 5 kV, spot size 2, 30 µm aperture, and dwell time of 20 µs for a 3072 x 2048 image frame for biopsies. For the Zeiss Gemini, tiled images were acquired with ATLAS 5 software using a Sense BSD detector at an accelerating voltage of 5.0 kV reduced to a landing energy of 3 kV via a 2 kV Tandem Decel stage bias, at a working distance of 5.082 mm, with a current of 400 pA, and dwell time of 3.2 µs for a 8192 x 8192 image frames.

### Design and build of the tomoSTORM light microscope

The tomoSTORM light microscope, based on the modular low-cost open-source *openFrame* microscope stand, was designed to image serial ultrathin easyIRF sections in RF, WF and SMLM modes.

tomoSTORM was built on an *openFrame* base layer (OF-LL-BP) with M6 screw mounting holes to attach the frame to an optical bench or breadboard. Above the base layer was the detection layer (OF-LL-65) containing a 100% reflective 45 degree mirror to direct the reflected or fluorescence signal to the imaging camera. The TE-cooled CellCam Kikker 100MT CMOS camera was used, with a 0.35X demagnifier adapter (Motic, #1101001904111) positioned in front of the camera to reduce the image size to a third of the camera sensor. This enabled fewer sensor rows to be read out per image and hence increased the CMOS camera frame rate to be fast enough (> 33 fps) for practical SMLM acquisitions. The tube lens (Olympus 180mm tube lens, SWTLU-C) was mounted vertically on the side of the *openFrame* detection layer, positioned before the Motic adapter.

The *openFrame* epi illumination layer (Cairn Research Ltd, DCG/UNI/EMA) sat directly above the detection layer. This allowed manual switching between RF and WF modes. The RF cube incorporated a 10:90 R:T beamsplitter (Thorlabs BSN10R) and a neutral density ND4 filter (Thorlabs ND40B) positioned in the excitation filter port of the cube to limit the excitation laser power to avoid photobleaching the sample. The reflection contrast was due to the refractive index difference at the sample/substrate and sample/air interface. A multiband fluorescent cube included a dichroic beamsplitter for excitation laser lines at 405 nm, 465 nm and 635 nm (Chroma ZT405/465/625/825rpc-UF2) and complementary multiband excitation (Chroma ZET405/465/625/825x) and emission filters (Chroma ZET405/465/625/825m).

The optical fibre-coupled excitation source entered through the side port of the epi-illumination layer via a critical illumination unit. The multiline multimode diode laserbank provided excitation at 405 nm (0.35 W), 465 nm (3.5 W), 520 nm (1 W) and 635 nm (0.7 W) via a single multimode optical fibre (Thorlabs M59L02 1000 um 0.5 NA). This optical fibre was vibrated using a de-speckler unit (Cairn Research Ltd) to mix the propagating optical modes, and was butt-coupled to a 20 cm long square core fibre (OBS Fiber, 1000 μm core, 0.22 NA). The square core optical fibre further enhanced the mixing of the optical modes to provide approximately uniform power density via critical illumination across the ∼70 μm x 70 μm FOV, and therefore approximately uniform achievable resolution. The critical illumination module included a zoom lens to image the square core of the fibre onto an adjustable rectangular aperture to maximise the power transmitted to the sample. This aperture was imaged onto the sample to enable accurate tessellation for tiled imaging with minimal photobleaching outside the square FOV that was set to match the observable FOV on the rectangular CMOS sensor.

The focus layer (Cairn Research Ltd, P1500/OBJ/NEC) incorporated a stepper motor-based z-stage (Cairn Z-Act) that supported a manually adjustable objective lens turret for 4 RMS thread objectives (Thorlabs OT1). The objective lenses used were a 10x/ 0.3 NA objective lens (Amscope PF10X-INF) and a 100x/ 1.3 NA objective lens (Amscope PF100X-INF). The focus layer also contained a 775 nm short pass dichroic beamsplitter (Chroma ZT775sp-2p) to introduce infrared radiation from a superluminescent diode (SLD) source (Lightley et al., 2023) which was focused onto the microscope sample plane to provide an autofocus signal. The autofocus was used to continuously correct for drift during SMLM acquisitions.

The stage adapter layer sat above the focus layer, and supported an encoded motorised microscope xy stage (Märzhäuser Wetzlar GmbH, SCAN IM 130 x 85) used for image tiling and moving between selected ROIs in sequential serial sections. A Märzhäuser TANGO 2 DT controller was used to control the xy stage and interface to the 2-axis joystick. A transillumination pillar was positioned above the sample supporting a white LED source (Cairn MonoLED).

Micro-Magellan was used as a pre-installed acquisition tool in Micro-Manager to acquire the tiled images (Pinkard et al., 2016). The autofocus was run as a Micro-Manager 2.0 (Edelstein et al., 2014) autofocus plugin (https://github.com/ImperialCollegeLondon/openAF). The multi-dimensional acquisition (MDA) module in Micro-Manager 2.0 was used to control all hardware during SMLM acquisition.

### TomoSTORM imaging in RF and WF modes

Serial ultrathin easyIRF sections were imaged with the 10x/ 0.3 NA Amscope objective lens in both RF and WF modes. The sample was critically illuminated such that the rectangular mask was imaged onto the sample. The FOV of the camera sensor was cropped to match the size of the image of the mask. The pixel size was calibrated using the automatic pixel calibration tool in Micro-Manager and a USAF 1951 resolution test chart illuminated in trans-illumination mode.

Once the pixel size for the system had been calibrated, the ultrathin sections could be imaged. Sections were imaged in RF mode first by manually switching to the reflection dichroic cube. The sample was imaged first at low excitation power density (3 μW/cm^2^) with an exposure of 50 ms. The explore functionality within the MicroMagellan plugin was used to identify the area of the coverslip containing the sections to be tiled. A 0% overlap between the tiles was used. A tiled image was built up progressively within the MicroMagellan graphical user interface (GUI) by highlighting regions and confirming acquisition of those tiles, after which the stage automatically moved in a snake-like pattern to acquire images. Once the sections had been explored, tiles of interest were selected using the grid and surfaces tab within MicroMagellan. The number of rows and columns of the grid was increased to cover the imaging region with the required number of tiles. The xy positions of the resulting grid were exported to Micro-Manager’s position list using the MicroMagellan main panel. With the sample in focus, a stack of ome-tiff images were acquired from the defined stage positions in the position list using the MDA.

The imaging modality was then switched to WF by manually switching to the multiband fluorescence dichroic. If the image of the mask occurred at a different lateral position on the camera sensor, the cropped FOV on the camera was redefined and matched in size to the FOV defined for the RF tiles. The same procedure was followed to acquire the tiled WF images with a power density (0.375 W/cm^2^) and an exposure of 200 ms.

The low magnification image tiles were stitched to create an RF montage and a WF montage using an ImageJ macro in Fiji. The RF and WF maps were then overlaid. To overlay, the RF and WF images were first imported into the BigWarp plugin in Fiji. The WF image was designated as the ‘moving image’ and RF image as the ‘target image’. Four landmarks were added to the corners of the serial sections in both the target image and the moving image. A translation transformation was used to bring the images into alignment using the merge function in Fiji. The resulting overlaid RF/WF image was used to support navigation between sections in the tomoSTORM microscope, to select ROIs for higher magnification LM imaging, and to target sections and ROIs for downstream SMLM and EM imaging.

The 100x/ 1.3 NA oil objective lens was then used to image one section in WF fluorescence mode to help select ROIs for subsequent imaging of the whole cell volume in SMLM mode across sequential sections, with a WF image of each fluorescence ROI being recorded prior to SMLM acquisition. The WF images were usually recorded using a 200 ms exposure.

### Power titration for imaging in SMLM mode on the tomoSTORM microscope

A power titration experiment was undertaken to determine the optimal power density for SMLM of GFP (Supp.Fig.1) and AF488 (Supp.Fig.2) in easyIRF sections to ensure sufficient fluorophore blinking while minimizing photobleaching. To avoid photobleaching artefacts due to repeated measurements, different sections with similar characteristics on the same substrate were measured at varying power levels. Each measurement was analysed using the standard localisation pipeline, extracting the average intensity and visually comparing it to individual raw frames (Supp.Fig.1a & 2a) to assess the signal-to-noise ratio (SNR) and determine whether most fluorophores were successfully driven into the dark state. The background fluorescence and bleaching characteristics at different power densities were also analysed (Supp.Fig.1b,c & 2b,c). The optimal power density was selected based on achieving a strong signal for reliable localisation while sending non-target fluorophores into the dark state and keeping a reasonable background level. Lastly, blinking dynamics were assessed by tracking localisation events over time to determine the average fluorophore emission duration, which consistently measured approximately two frames (Supp.Fig.1d & 2d). Thus, HeLa cells and hIPSC-derived cortical neurons expressing mitoGFP were imaged in SMLM mode using a 330W/cm² power density at the sample at 465 nm excitation, and the hIPSC-derived cortical neurons containing AF488 were imaged with a power density of 1231 W/cm² at 465 nm excitation.

### SMLM acquisition on the tomoSTORM

For each SMLM acquisition, 10,000 frames were acquired at 30 ms exposure per frame. A power density of 330 W/cm^2^ at the sample plane was typically used to drive blinking of GFP and a power density of 1231 W/cm^2^ for the Alexa dyes. The autofocus was set to provide continuous focal correction during the SMLM acquisition. The excitation power was set to induce blinking and switching of fluorophores to the dark state. Once an adequate proportion of the fluorophore population was in the dark state, such that fluorescent signal from single neighbouring emitters was not overlapping, the acquisition of the SMLM frames was started using the MDA. The fluorophores were typically excited for approximately 10 s to switch sufficient fluorophores into the dark state. Once an SMLM acquisition of an ROI was complete, the stage was moved to image either a separate ROI within the same section or to the same ROI in the following section.

### TomoSTORM SMLM reconstruction

The raw fluorescence images acquired using tomoSTORM were processed using the Picasso software package to generate SMLM reconstructions. Single molecule localisation and fitting were performed using Picasso Localize with settings optimised individually for each dataset. The box side length for single molecule fitting was set to be larger than three times the typical PSF size (11 pixels in this case), and the minimum net gradient was adjusted (typically between 1500 and 2000) to maximize the number of valid detections while minimizing false positives from background noise. Drift correction was applied using redundant cross-correlation, dividing the dataset into 10 segments, as implemented in Picasso Render.

The localised data were further filtered using a custom Python script, which relies on the Picasso and Nanopyx (Saraiva et al., 2025) libraries. Specifically, localisations were filtered based on photon count, sigma values, background intensity, and density. The lower photon threshold was set at the 1st percentile of the average photon count of the stack of frames, ensuring the retention of only the brightest signals without imposing an upper limit. Sigma values were estimated by calculating the expected full width at half maximum (FWHM) and deriving the corresponding Gaussian width. A lower threshold was then set at 80% of this value, while the upper limit remained unrestricted. Background intensity filtering played a crucial role in rejecting spurious detections. The acceptance range was defined between one standard deviation below the mean background and two standard deviations above it, effectively minimizing background noise while preserving localisations corresponding to structural features. Finally, a density filter was applied to further reduce background noise. Localisations were required to have at least 20 neighbouring detections within a 500 nm radius to be considered valid. Any localisations with fewer neighbours were discarded.

Following data refinement, the super-resolved images were rendered at the desired magnification, typically 5x in this study. The resolution of the final reconstructed dataset was quantified using decorrelation analysis, providing an accurate measurement of the achieved resolution.

### Design of a napari interface for section and ROI selection

To target sections and ROIs imaged by LM for downstream EM, a napari plugin was designed and built, named napari-mass (for Microscopy Array Section Setup; MASS). The plugin allowed the user to: import the overlaid RF/WF image; annotate one of the sections using the napari drawing tools (specifically the polygon tool); copy and paste the annotation to each subsequent section; view the annotated sections as a stack in the template viewer; and select and propagate fluorescent ROIs through the stack of serial sections. The napari-mass plugin allowed configuration of parameters such as the location of the image(s), which were stored in a project file for convenience. Additionally, all section and ROI locations were stored in an output file for downstream use in targeted EM imaging. The napari-mass plugin was written in Python and used the napari-ome-zarr, ome-zarr, and tifffile packages to support OME file formats. The napari-mass plugin also used the dask-image, scikit and OpenCV packages for image processing. The ROI propagation and template view functions were inspired by the propagation and local view concepts (respectively) of the MagFinder Fiji plugin (https://github.com/templiert/MagFinder).

### Post-staining sections to add electron contrast

After imaging with LM, the coverslip was removed from the slide holder and cleaned with ethanol to remove oil. The sections were baked at 60° for 10 mins on a hot plate to facilitate attachment (section-side-up). Sections were post-stained with a combination of 2% uranyl acetate in ddH_2_O (w/v) with 0.5% potassium permanganate for 10 mins at room temperature in the dark. Coverslips were then floated on a droplet of water for 30 secs (section-side-down) followed by dipping the coverslip 10 times in each of five beakers of water. Sections were then stained in Reynold’s lead citrate for 10 min and washed as before. Coverslips were air dried, mounted onto an SEM stub with a carbon sticky tab and the edges coated with silver paint on the sides to improve conductivity.

### Implementing array tomography on the Phenom Pharos tabletop FEG SEM

The tabletop Phenom Pharos G2 FEG SEM (Thermo Fisher Scientific) was used to image ultrathin easyIRF sections. The open-source software SBEMimage (Titze et al., 2018) was used to control array tomography through the Phenom Pharos Application Programming Interface (API; Thermo Fisher Scientific). A new module called ‘array’ was designed and implemented within SBEMimage for this purpose. The module builds and improves upon the existing MagC module of SBEMimage (Collectome LLC, Switzerland).

An overview image of the stub was acquired using the NavCAM to find the position of the section ribbon on the coverslip. The Pharos was then switched to electron imaging mode. The NavCAM image was used to guide placement of a SBEMimage ‘grid 0’ to guide acquisition of an electron image of the section ribbon at 500 nm pixel size and 0.2 μs dwell time. The merged RF/WF images and the array data file containing xy coordinates of fluorescent ROIs defined in napari-mass were imported into SBEMimage using the array module. The imported RF/WF image was aligned to the EM image of the section ribbon. This alignment was used as a calibration to transfer the ROI positions into the EM stage coordinate system, enabling the user to find and selectively image the ROIs. For each ROI, the working distance (focus) was manually adjusted, and then an image was acquired quickly at 10 or 20 nm pixel size and 0.1 μs dwell time to ensure that the ROI was present in the image with minimal mass loss due to beam-section interaction. Once the ROI was confirmed, higher quality images were acquired at 20 and 10 nm with dwell times of 5 and 10 μs respectively. For all ROIs, high resolution images were acquired using the BSE detector with imaging parameters set to 3 kV, with a spot size of 3.9.

### EM image processing: Manual lateral tile stitching

EM image tiles were manually stitched in Fiji using the TrakEM2 plugin. Individual tiles were imported into a single TrakEM2 layer corresponding to the section of interest. The tiles were initially arranged using metadata when available, and manual adjustments were performed to refine their relative positions. Stitching was guided by overlapping structural features such as membrane contours, nuclei, and organelles to ensure continuity across tile boundaries. When necessary, transparency and contrast settings were adjusted to facilitate visual alignment. Once satisfactory stitching was achieved, the layer was flattened into a composite image, which was then exported for subsequent axial alignment and overlay.

### EM Image processing: Manual axial image registration

Stitched EM images were aligned along the z-axis using the TrakEM2 plugin in Fiji. The process involved applying rigid transformations (translations and rotations) to ensure continuity and accurate structural correspondence across serial sections. Images were imported into separate layers within a single TrakEM2 project, and the layers were unlinked to allow individual manipulation. An initial rough alignment was performed using the automatic affine alignment tool. This was followed by manual refinement, where each subsequent section was adjusted in the x and y axes and rotated in-plane based on visual inspection of consistent ultrastructural features such as organelles and membranes. The stack was iteratively refined to produce a continuous and coherent 3D reconstruction, prioritizing the preservation of ultrastructural integrity. The final aligned dataset was then exported for downstream integration into the 3D volume. For this pipeline, only the EM images were aligned axially. However, should axial alignment of the LM images be required, the same approach could be used.

### EM image processing: Automated tile stitching

A registration pipeline was developed named ‘muvis-align’ to facilitate automated lateral and axial tile stitching. The pipeline was capable of handling large image datasets in the GB and TB regime, and included a number of pre- and post-processing steps. The pipeline was written in Python and based on a number of open-source Python packages, with the main base package building on multiview-stitcher [https://doi.org/10.5281/zenodo.13151252], a modular toolbox for distributed and tiled stitching. The registration method used phase correlation to determine the optimal image alignment. The pipeline leveraged the latest Open Microscopy Environment (OME) Next Generation File Formats (NGFF) (Moore et al., 2021). The main Python packages underpinning the pipeline were dask, xarray, spatial-image, zarr [https://doi.org/10.5281/zenodo.3773450] and ome-zarr.

### Correlative image processing: Overlay of LM and EM images

WF images were registered to their corresponding EM sections using the BigWarp plugin in Fiji. For each section, the WF image was designated as the ‘moving’ image and the EM image as the ‘target’. Prominent ultrastructural features visible in both modalities were selected as landmarks to compute the transformation. In cases where few such features were apparent, contrast enhancement of the WF image revealed cell structure-associated autofluorescence, effectively outlining cell contours that could be matched to their EM counterparts. After the initial overlay, landmarks were refined by zooming into specific features to improve local alignment until the overlay was satisfactory. The transformed WF image was then exported at the same resolution as the EM image. Once all sections were accurately overlaid, they were stacked to reconstruct the complete 3D dataset.

Registration of the SMLM images to the EM images was challenging even for the benchmark samples, where mitochondria were visible in LM (mitoGFP) and EM (recognisable morphology). The challenge was even greater for the experimental system (hIPSC-derived cortical neurons with ɑ-Syn-AF488), where most localisations were punctate and the underlying ultrastructure of the target cell compartments was unknown. In these cases, WF images served as an effective intermediate step to facilitate the registration of SMLM onto EM where direct registration was not feasible. Once the WF stack had been registered to the EM stack, the corresponding SMLM images were registered to the EM-registered WF images using BigWarp. By aligning SMLM to the EM-registered WF stack, a reliable initial registration of SMLM to EM was achieved, which could then be refined to generate a high-precision SMLM-EM overlay.

## Resources

dx.doi.org/10.17504/protocols.io.3byl498x8go5/v1

https://napari.org

https://github.com/SBEMimage

https://github.com/ImperialCollegeLondon/openAF

https://doi.org/10.5281/zenodo.13151252

https://doi.org/10.5281/zenodo.15029515

## Acknowledgements

Human samples used in this research project (reference R14094) were obtained from the Imperial College Healthcare Tissue Bank (ICHTB). ICHTB is supported by the National Institute for Health Research (NIHR) Biomedical Research Centre based at Imperial College Healthcare NHS Trust and Imperial College London. ICHTB is approved by Wales REC3 to release human material for research (22/WA/2836). The authors acknowledge and thank: Raffaella Carzaniga for supporting the early stages of the pipeline development; Pippa Hawes for critical reading of the manuscript; Katherine Lau and Guido Ridolfi from DELMIC B.V. for support with the specification and procurement of the Phenom Pharos and design of the coverslip holders; James Carr, Ben Bower and Tom Warwick from Blue Scientific for assistance with the specification and shipping of the Phenom Pharos; Wouter Arts, Keriya Mam and Dlangir Cordero from Thermo Fisher Scientific for API and software support and optimisation of the Phenom Pharos performance for imaging easyIRF sections; Thomas Templier for support and discussions in development of napari and SBEMimage plugins; Loren Looger, Maria G. Paez-Segala and Ben Campbell for discussions about fluorescent probes for IRF; and the Crick Grants, Finance and Legal teams and the Stellenbosch University Legal team for support during the course of the project and in the shipping of the VP-CLEM-Kit to South Africa.

## Funding acknowledgements

The authors thank the Chan Zuckerberg Initiative for funding through the Visual Proteomics Imaging programme (no. vpi-0000000044; https://doi.org/10.37921/743590vtudfp) awarded to Collinson, French and Henriques. This work was supported by the Francis Crick Institute which receives its core funding from Cancer Research UK (CC1076), the UK Medical Research Council (CC1076), and the Wellcome Trust (CC1076), and through the Crick Idea to Innovation (i2i) scheme. R. Henriques and A. G. Vesga express their gratitude for the support of the Gulbenkian Foundation (Fundação Calouste Gulbenkian); the European Research Council under the European Union’s Horizon 2020 research and innovation programme (grant agreement no. 101001332 and Marie Skłodowska-Curie grant agreement no. 101180631); and the European Molecular Biology Organization Installation Grant (no. EMBO-2020-IG-4734). R.Henriques acknowledges funding from the Fundação para a Ciência e Tecnologia (Portugal) through the MOSTMICRO-ITQB R&D Unit (nos. UIDB/04612/2020 and UIDP/04612/2020).

---

**Supplementary Figure 1.**
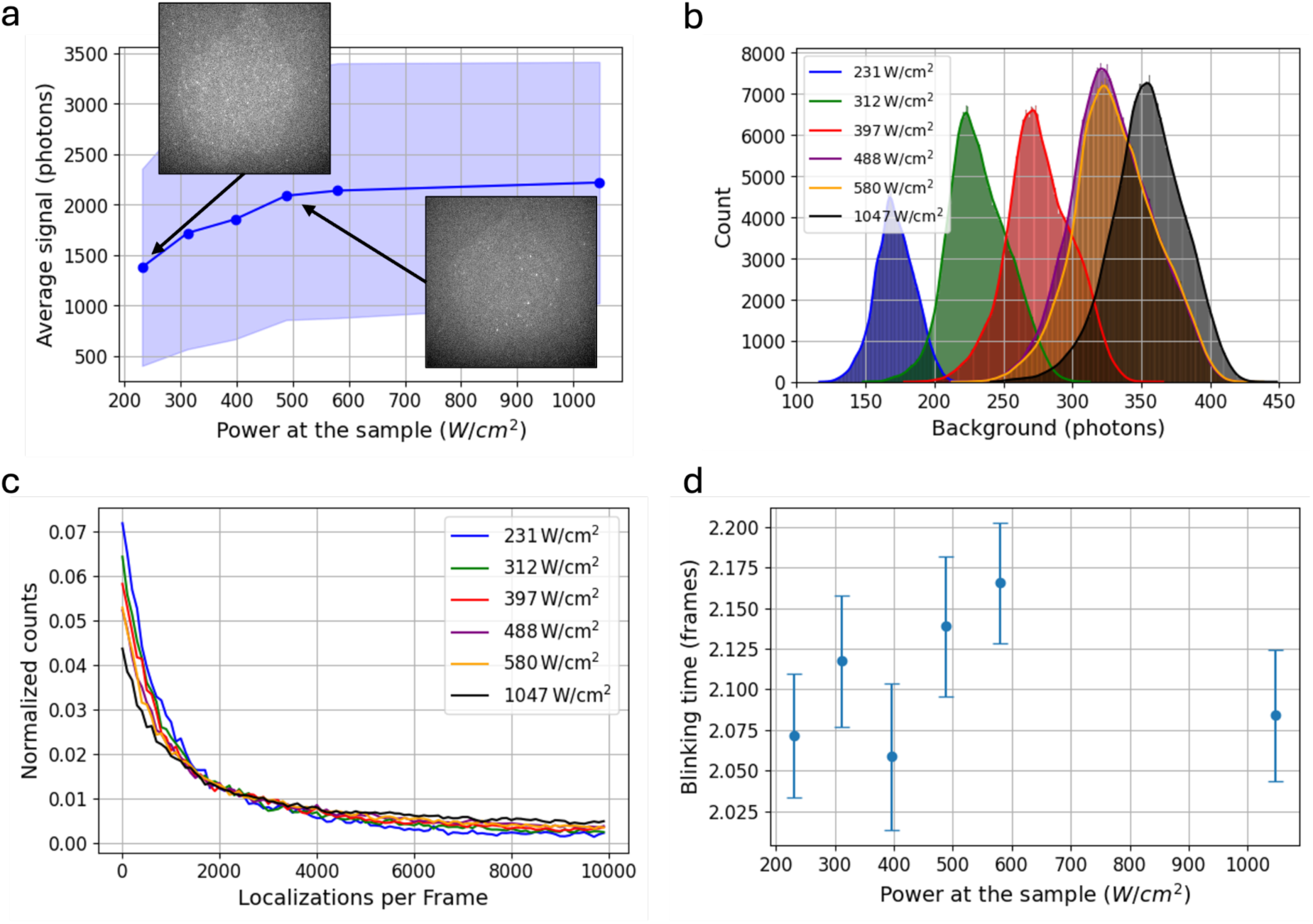
Optimization of illumination power for SMLM in easyIRF sections containing HeLa cells expressing mitoGFP. (a) Power titration curve showing the average signal per frame as a function of illumination power of the same FOV, but for different sections of the same cell for each power level. At lower illumination powers, not all fluorophores transition to the dark state, leading to high background and suboptimal conditions for SMLM, as seen in the inset images. Increasing power effectively drives fluorophores into the dark state, reducing background fluorescence. (b) Histogram of background photon counts for different illumination powers, demonstrating a progressive increase in background signal with increasing power. (c) Bleaching curves normalized to total background signal, showing the total number of localisations per frame for different illumination powers. Strong bleaching is observed at both low and high powers. (d) Relationship between power and the average blinking time per fluorophore. No significant differences in blinking time were observed across different power densities. Consequently, a power density of 330 W/cm² was selected as it provided the best balance between minimizing background fluorescence and maintaining a strong SNR.

**Supplementary Figure 2.**
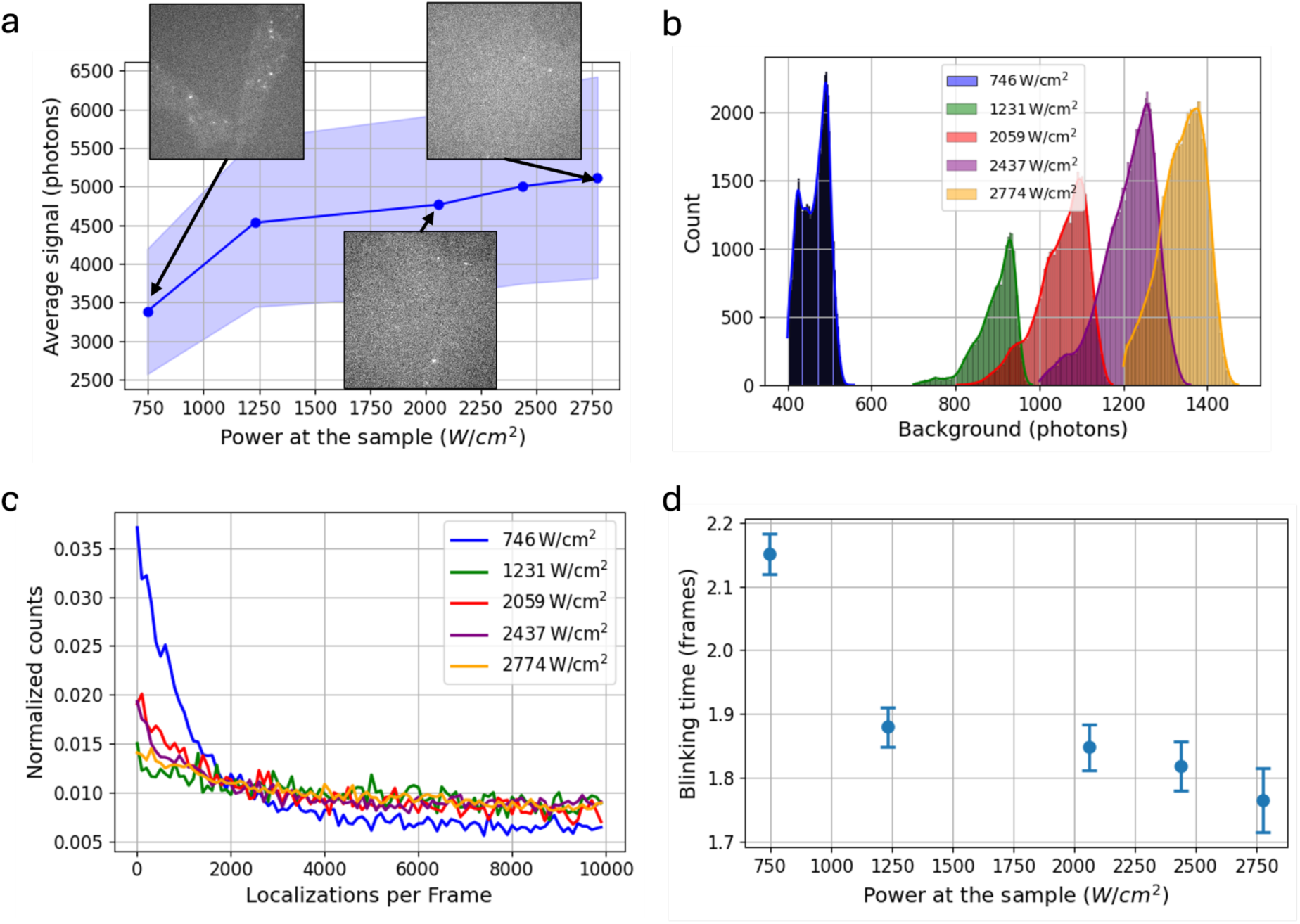
Optimization of illumination power for SMLM in easyIRF sections containing hiPSC-derived cortical neurons exposed to ɑ-Syn-AF488. (a) Power titration curve showing the average signal per frame as a function of illumination power. At lower illumination powers, not all fluorophores transition to the dark state, leading to high background and suboptimal conditions for SMLM, as seen in the inset images. Increasing power effectively drives fluorophores into the dark state, reducing background fluorescence. (b) Histogram of background photon counts for different illumination powers, demonstrating a progressive increase in background fluorescence with increasing power. (c) Bleaching curves normalized to the area of the background histograms, showing the total localisation counts per frame for different illumination powers. Strong bleaching is observed at both low and high powers. (d) Relationship between power and the average blinking time per fluorophore, indicating that 1231 W/cm² provides the best compromise, with a low background, efficient dark-state transition, and minimal reduction in blinking time.

## References

Baatsen, P., S. Gabarre, K. Vints, R. Wouters, D. Vandael, R. Goodchild, S. Munck, and N.V. Gounko. 2021. Preservation of Fluorescence Signal and Imaging Optimization for Integrated Light and Electron Microscopy. Frontiers in Cell and Developmental Biology. 9.

Bogovic, J.A., P. Hanslovsky, A. Wong, and S. Saalfeld. 2016. Robust registration of calcium images by learned contrast synthesis. In 2016 IEEE 13th International Symposium on Biomedical Imaging (ISBI). 1123-1126.

Briggs, J.A. 2013. Structural biology in situ--the potential of subtomogram averaging. Current opinion in structural biology. 23:261–267.

Campbell, B.C., M.G. Paez-Segala, L.L. Looger, G.A. Petsko, and C.F. Liu. 2022. Chemically stable fluorescent proteins for advanced microscopy. Nature methods. 19:1612–1621.

Cardona, A., S. Saalfeld, J. Schindelin, I. Arganda-Carreras, S. Preibisch, M. Longair, P. Tomancak, V. Hartenstein, and R.J. Douglas. 2012. TrakEM2 software for neural circuit reconstruction. PLoS One. 7:e38011.

Chiu, C.-L., N. Clack, and t.n. community. 2022. napari: a Python Multi-Dimensional Image Viewer Platform for the Research Community. Microscopy and Microanalysis. 28:1576–1577.

Choi, M.L., A. Chappard, B.P. Singh, C. Maclachlan, M. Rodrigues, E.I. Fedotova, A.V. Berezhnov, S. De, C.J. Peddie, D. Athauda, G.S. Virdi, W. Zhang, J.R. Evans, A.I. Wernick, Z.S. Zanjani, P.R. Angelova, N. Esteras, A.Y. Vinokurov, K. Morris, K. Jeacock, L. Tosatto, D. Little, P. Gissen, D.J. Clarke, T. Kunath, L. Collinson, D. Klenerman, A.Y. Abramov, M.H. Horrocks, and S. Gandhi. 2022. Pathological structural conversion of α-synuclein at the mitochondria induces neuronal toxicity. Nat Neurosci. 25:1134–1148.

Culley, S., D. Albrecht, C. Jacobs, P.M. Pereira, C. Leterrier, J. Mercer, and R. Henriques. 2018. Quantitative mapping and minimization of super-resolution optical imaging artifacts. Nature methods. 15:263–266.

Denk, W., and H. Horstmann. 2004. Serial block-face scanning electron microscopy to reconstruct three-dimensional tissue nanostructure. PLoS Biol. 2:e329.

Descloux, A., K.S. Grußmayer, and A. Radenovic. 2019. Parameter-free image resolution estimation based on decorrelation analysis. Nature methods. 16:918–924.

Edelstein, A.D., M.A. Tsuchida, N. Amodaj, H. Pinkard, R.D. Vale, and N. Stuurman. 2014. Advanced methods of microscope control using muManager software. Journal of biological methods. 1.

Fu, Z., D. Peng, M. Zhang, F. Xue, R. Zhang, W. He, T. Xu, and P. Xu. 2020. mEosEM withstands osmium staining and Epon embedding for super-resolution CLEM. Nature methods. 17:55–58.

Fulton, K.A., P.V. Watkins, and K.L. Briggman. 2024. GAUSS-EM, guided accumulation of ultrathin serial sections with a static magnetic field for volume electron microscopy. Cell Rep Methods. 4:100720.

Genoud, C., G.W. Knott, K. Sakata, B. Lu, and E. Welker. 2004. Altered synapse formation in the adult somatosensory cortex of brain-derived neurotrophic factor heterozygote mice. The Journal of neuroscience : the official journal of the Society for Neuroscience. 24:2394–2400.

Graham, B.J., D.G.C. Hildebrand, A.T. Kuan, J.T. Maniates-Selvin, L.A. Thomas, B.L. Shanny, and W.-C.A. Lee. 2019. High-throughput transmission electron microscopy with automated serial sectioning. bioRxiv:657346.

Guerin, C.J., and S. Lippens. 2021. Correlative light and volume electron microscopy (vCLEM): How community participation can advance developing technologies. J Microsc. 284:97–102.

Harris, K., F. Jensen, and B. Tsao. 1992. Three-dimensional structure of dendritic spines and synapses in rat hippocampus (CA1) at postnatal day 15 and adult ages: implications for the maturation of synaptic physiology and long-term potentiation [published erratum appears in J Neurosci 1992 Aug;12(8):following table of contents]. The Journal of Neuroscience. 12:2685–2705.

Hayworth, K., N. Kasthuri, R. Schalek, and J. Lichtman. 2006. Automating the Collection of Ultrathin Serial Sections for Large Volume TEM Reconstructions. Microscopy and Microanalysis. 12:86–87.

Heiligenstein, X., and M.S. Lucas. 2022. One for All, All for One: A Close Look at In-Resin Fluorescence Protocols for CLEM. Front Cell Dev Biol. 10:866472.

Hoyer, W., T. Antony, D. Cherny, G. Heim, T.M. Jovin, and V. Subramaniam. 2002. Dependence of alpha-synuclein aggregate morphology on solution conditions. J Mol Biol. 322:383–393.

Hutchings, J., and E. Villa. 2024. Expanding insights from in situ cryo-EM. Current opinion in structural biology. 88:102885.

Ignatiou, A., K. Macé, A. Redzej, T.R.D. Costa, G. Waksman, and E.V. Orlova. 2024. Structural Analysis of Protein Complexes by Cryo-Electron Microscopy. Methods Mol Biol. 2715:431–470.

Johnson, E., and R. Kaufmann. 2017. Correlative In-Resin Super-Resolution Fluorescence and Electron Microscopy of Cultured Cells. Methods Mol Biol. 1663:163–177.

Knott, G., H. Marchman, D. Wall, and B. Lich. 2008. Serial section scanning electron microscopy of adult brain tissue using focused ion beam milling. The Journal of neuroscience : the official journal of the Society for Neuroscience. 28:2959–2964.

Krentzel, D., M. Elphick, M.-C. Domart, C.J. Peddie, R.F. Laine, C. Shand, R. Henriques, L.M. Collinson, and M.L. Jones. 2024. CLEM-Reg: An automated point cloud based registration algorithm for correlative light and volume electron microscopy. bioRxiv.

Kukulski, W., M. Schorb, S. Welsch, A. Picco, M. Kaksonen, and J.A. Briggs. 2011. Correlated fluorescence and 3D electron microscopy with high sensitivity and spatial precision. The Journal of cell biology. 192:111–119.

Kwakwa, K., A. Savell, T. Davies, I. Munro, S. Parrinello, M.A. Purbhoo, C. Dunsby, M.A. Neil, and P.M. French. 2016. easySTORM: a robust, lower-cost approach to localisation and TIRF microscopy. Journal of biophotonics. 9:948–957.

Li, X., G. Ji, X. Chen, W. Ding, L. Sun, W. Xu, H. Han, and F. Sun. 2017. Large scale three-dimensional reconstruction of an entire Caenorhabditis elegans larva using AutoCUTS-SEM. Journal of structural biology. 200:87–96.

Lightley, J., S. Kumar, M.Q. Lim, E. Garcia, F. Görlitz, Y. Alexandrov, T. Parrado, C. Hollick, E. Steele, K. Roßmann, J. Graham, J. Broichhagen, I.A. McNeish, C.A. Roufosse, M.A.A. Neil, C. Dunsby, and P.M.W. French. 2023. openFrame: A modular, sustainable, open microscopy platform with single-shot, dual-axis optical autofocus module providing high precision and long range of operation. J Microsc. 292:64–77.

McCafferty, C.L., S. Klumpe, R.E. Amaro, W. Kukulski, L. Collinson, and B.D. Engel. 2024. Integrating cellular electron microscopy with multimodal data to explore biology across space and time. Cell. 187:563–584.

Micheva, K.D., B. Gong, F. Collman, R.J. Weinberg, S.J. Smith, J.S. Trimmer, and K.D. Murray. 2023. Developing a Toolbox of Antibodies Validated for Array Tomography-Based Imaging of Brain Synapses. Eneuro. 10.

Micheva, K.D., and S.J. Smith. 2007. Array tomography: a new tool for imaging the molecular architecture and ultrastructure of neural circuits. Neuron. 55:25–36.

Moore, J., C. Allan, S. Besson, J.M. Burel, E. Diel, D. Gault, K. Kozlowski, D. Lindner, M. Linkert, T. Manz, W. Moore, C. Pape, C. Tischer, and J.R. Swedlow. 2021. OME-NGFF: a next-generation file format for expanding bioimaging data-access strategies. Nature methods. 18:1496–1498.

Nickell, S., C. Kofler, A.P. Leis, and W. Baumeister. 2006. A visual approach to proteomics. Nature reviews. Molecular cell biology. 7:225–230.

Paez-Segala, M.G., M.G. Sun, G. Shtengel, S. Viswanathan, M.A. Baird, J.J. Macklin, R. Patel, J.R. Allen, E.S. Howe, G. Piszczek, H.F. Hess, M.W. Davidson, Y. Wang, and L.L. Looger. 2015. Fixation-resistant photoactivatable fluorescent proteins for CLEM. Nature methods. 12:215–218, 214 p following 218.

Peddie, C.J., K. Blight, E. Wilson, C. Melia, J. Marrison, R. Carzaniga, M.C. Domart, P. O’Toole, B. Larijani, and L.M. Collinson. 2014. Correlative and integrated light and electron microscopy of in-resin GFP fluorescence, used to localise diacylglycerol in mammalian cells. Ultramicroscopy. 143:3–14.

Peddie, C.J., M.-C. Domart, X. Snetkov, P. O’Toole, B. Larijani, M. Way, S. Cox, and L.M. Collinson. 2017. Correlative super-resolution fluorescence and electron microscopy using conventional fluorescent proteins in vacuo. Journal of structural biology. 199:120–131.

Peddie, C.J., C. Genoud, A. Kreshuk, K. Meechan, K.D. Micheva, K. Narayan, C. Pape, R.G. Parton, N.L. Schieber, Y. Schwab, B. Titze, P. Verkade, A. Weigel, and L.M. Collinson. 2022. Volume electron microscopy. Nat Rev Methods Primers. 2:51.

Peng, D., N. Li, W. He, K.R. Drasbek, T. Xu, M. Zhang, and P. Xu. 2022. Improved Fluorescent Proteins for Dual-Colour Post-Embedding CLEM. Cells. 11:1077.

Pinkard, H., N. Stuurman, K. Corbin, R. Vale, and M.F. Krummel. 2016. Micro-Magellan: open-source, sample-adaptive, acquisition software for optical microscopy. Nature methods. 13:807–809.

Polishchuk, R.S., E.V. Polishchuk, P. Marra, S. Alberti, R. Buccione, A. Luini, and A.A. Mironov. 2000. Correlative light-electron microscopy reveals the tubular-saccular ultrastructure of carriers operating between Golgi apparatus and plasma membrane. The Journal of cell biology. 148:45–58.

Sanada, T., J. Yamaguchi, Y. Furuta, S. Kakuta, I. Tanida, and Y. Uchiyama. 2022. In-resin CLEM of Epon-embedded cells using proximity labeling. Scientific reports. 12:11130.

Saraiva, B.M., I. Cunha, A.D. Brito, G. Follain, R. Portela, R. Haase, P.M. Pereira, G. Jacquemet, and R. Henriques. 2025. Efficiently accelerated bioimage analysis with NanoPyx, a Liquid Engine-powered Python framework. Nature methods. 22:283–286.

Schindelin, J., C.T. Rueden, M.C. Hiner, and K.W. Eliceiri. 2015. The ImageJ ecosystem: An open platform for biomedical image analysis. Molecular Reproduction and Development. 82:518–529.

Schnitzbauer, J., M.T. Strauss, T. Schlichthaerle, F. Schueder, and R. Jungmann. 2017. Super-resolution microscopy with DNA-PAINT. Nat Protoc. 12:1198–1228.

Sergey, G., K. Denis, H. Ava, G. Gediminas, O. Viola, K.O. L, L.R. H.P, B.M. O’, P. Roger, W.J. C, and d.M. Alex. 2018. Oxygen plasma focused ion beam scanning electron microscopy for biological samples. bioRxiv:457820.

Shi, Y., P. Kirwan, and F.J. Livesey. 2012. Directed differentiation of human pluripotent stem cells to cerebral cortex neurons and neural networks. Nat Protoc. 7:1836–1846.

Soto, G.E., S.J. Young, M.E. Martone, T.J. Deerinck, S. Lamont, B.O. Carragher, K. Hama, and M.H. Ellisman. 1994. Serial section electron tomography: a method for three-dimensional reconstruction of large structures. NeuroImage. 1:230–243.

Tanida, I., Y. Furuta, J. Yamaguchi, S. Kakuta, J.A. Oliva Trejo, and Y. Uchiyama. 2020a. Two-color in-resin CLEM of Epon-embedded cells using osmium resistant green and red fluorescent proteins. Scientific reports. 10:21871.

Tanida, I., S. Kakuta, J.A. Oliva Trejo, and Y. Uchiyama. 2020b. Visualization of cytoplasmic organelles via in-resin CLEM using an osmium-resistant far-red protein. Scientific reports. 10:11314.

Tanida, I., J. Yamaguchi, C. Suzuki, S. Kakuta, and Y. Uchiyama. 2023. Application of immuno- and affinity labeling with fluorescent dyes to in-resin CLEM of Epon-embedded cells. Heliyon. 9:e17394.

Templier, T. 2019. MagC, magnetic collection of ultrathin sections for volumetric correlative light and electron microscopy. Elife. 8.

Thirunavukkuarasu, S., E.A. Jares-Erijman, and T.M. Jovin. 2008. Multiparametric fluorescence detection of early stages in the amyloid protein aggregation of pyrene-labeled alpha-synuclein. J Mol Biol. 378:1064–1073.

Titze, B., C. Genoud, and R.W. Friedrich. 2018. SBEMimage: Versatile Acquisition Control Software for Serial Block-Face Electron Microscopy. Frontiers in neural circuits. 12.

White, E.L., Y. Amitai, and M.J. Gutnick. 1994. A comparison of synapses onto the somata of intrinsically bursting and regular spiking neurons in layer V of rat SmI cortex. Journal of Comparative Neurology. 342:1–14.

White, I.J., and J.J. Burden. 2023. A practical guide to starting SEM array tomography-An accessible volume EM technique. Methods Cell Biol. 177:171–196.

White, J.G., E. Southgate, J.N. Thomson, and S. Brenner. 1986. The structure of the nervous system of the nematode Caenorhabditis elegans. Philosophical Transactions Royal Soc Lond B Biological Sci. 314:1–340.

Xu, C.S., K.J. Hayworth, Z. Lu, P. Grob, A.M. Hassan, J.G. García-Cerdán, K.K. Niyogi, E. Nogales, R.J. Weinberg, and H.F. Hess. 2017. Enhanced FIB-SEM systems for large-volume 3D imaging. eLife. 6:e25916.

Yin, W., D. Brittain, J. Borseth, M.E. Scott, D. Williams, J. Perkins, C.S. Own, M. Murfitt, R.M. Torres, D. Kapner, G. Mahalingam, A. Bleckert, D. Castelli, D. Reid, W.-C.A. Lee, B.J. Graham, M. Takeno, D.J. Bumbarger, C. Farrell, R.C. Reid, and N.M. da Costa. 2020. A petascale automated imaging pipeline for mapping neuronal circuits with high-throughput transmission electron microscopy. Nature communications. 11:4949.

